# Environmental radiation exposure at Chornobyl has not systematically affected the genomes or mutagen tolerance phenotypes of local worms

**DOI:** 10.1101/2023.05.28.542665

**Authors:** Sophia C. Tintori, Derin Çağlar, Patrick Ortiz, Ihor Chyzhevskyi, Timothy A. Mousseau, Matthew V. Rockman

## Abstract

The 1986 disaster at the Chornobyl Nuclear Power Plant transformed the surrounding region into the most radioactive landscape known on the planet. Questions remain regarding whether this sudden environmental shift selected for species, or even individuals within a species, that are naturally more resistant to radiation exposure.

We collected, cultured, and cryopreserved 298 wild nematodes isolates from areas varying in radioactivity within the Chornobyl Exclusion Zone. We sequenced and assembled genomes *de novo* for 20 *Oschieus tipulae* strains, analyzed their genomes for evidence of recent mutation acquisition in the field and saw no evidence of an association between mutation and radiation level at the sites of collection. Multigenerational exposure of each of these strains to several mutagens in the lab revealed that strains vary heritably in tolerance to each mutagen, but mutagen tolerance cannot be predicted based on the radiation levels at collection sites.

## INTRODUCTION

Mutagens are a ubiquitous component of our environment – endogenous and exogenous, naturally occurring and anthropogenic. Of the tens of thousands of DNA lesions formed in each nucleus each day, the vast majority are corrected by our DNA repair machinery without incident. Even so, it is well established that heritable genetic variants in DNA repair components can contribute to cancer, birth defects, premature aging, and neurological defects (*1, 2*) .

Nematodes are emerging as a powerful tool for studying DNA repair (*3–5*) and toxin responses (*6, 7*), as well as natural variation in these processes (*8, 9*). They can reliably be collected from a broad range of environments across the planet (*10–12*), the wealth of published genomes makes them amenable to bioinformatic and quantitative genetics analyses (*10, 13*), and their small size and the extensive toolkit developed to study them allow researchers to experimentally test hypotheses generated from the previous two types of study. In this study we apply these methods, as well as new ones we have developed, to nematodes from the Chornobyl Exclusion Zone (CEZ), the site of the largest nuclear disaster in history.

Areas of the exclusion zone easily accessible to wildlife reach tens of sieverts per year, an order of magnitude higher than typical levels astronauts are exposed to during space travel (Supplementary Figure 1). Though animals can be found in these high radiation areas, evidence suggests that the exclusion zone may act as a population sink, at least for some highly vagile species (*14*), making it difficult to say how far back in an animal’s lineage they might have been exposed to these levels of radiation.

In recent years, ecological studies in the CEZ have reported species with abnormal physiology (*15, 16*), putatively due to radiation-induced DNA damage. Meanwhile, genetic studies have shown mixed evidence for detectable signatures of radiation damage in the genomes of surviving animals (*17–23*). To be clear, a lack of a recognizable mutagenic signature, or a lack of detectable selection pressure on tolerance alleles, does not suggest a harm-free environment.

For example, it is both true that (*17*) revealed no increase in *de novo* germline mutation rates in exposed parents who conceived after the disaster, and also that an untold number developed cancer due to their exposure. Thirty-seven years later researchers still strive to understand the extent and the implications of the damage done to life in this area.

In our study we collected nematode worms from locations throughout the CEZ ranging in ambient radiation levels from innocuous to dangerous. We use isohermaphrodite lines descended from these isolates for complementary experiments – first interrogating the genomes for genetic evidence of exposure in recent history, then experimentally testing their relative mutagen tolerance in a controlled setting.

## RESULTS

### Rhabditids collected from Chornobyl substrates spanning more than three orders of magnitude in ambient radioactivity

During two collecting trips to the Chornobyl Exclusion Zone, in August and October of 2019, we collected 237 local substrates (124 fruit, 65 soil and leaf litter, 18 invertebrates, 15 compost, 13 riparian plant stems, 1 wolf fecal sample, and 1 mushroom), as well as 16 substrates (apple and tomato) we placed as bait and returned to collect days later. We recovered nematodes from 157 of these substrates, and extracted 748 nematode isolates from these to establish cultures on nematode growth medium with *E. coli* as food source. We successfully established cultures for 298 of these isolates, representing 118 of the substrates collected, and we cryopreserved them for subsequent analysis. Ambient radiation at collection sites ranged from 2 to 4,786 mSv/y, as measured by Geiger counter, and substrates ranged from 0 to 2,019 Bq/gram dry weight, as measured by gamma spectrometer (Figure 1A). Ambient radiation and substrate radioactivity were linearly related when separated by substrate type, with soil substrates averaging 30-100x higher radioactivity than fruit substrates from the same collection site (Supplementary Figure 2).

**Figure 1:**
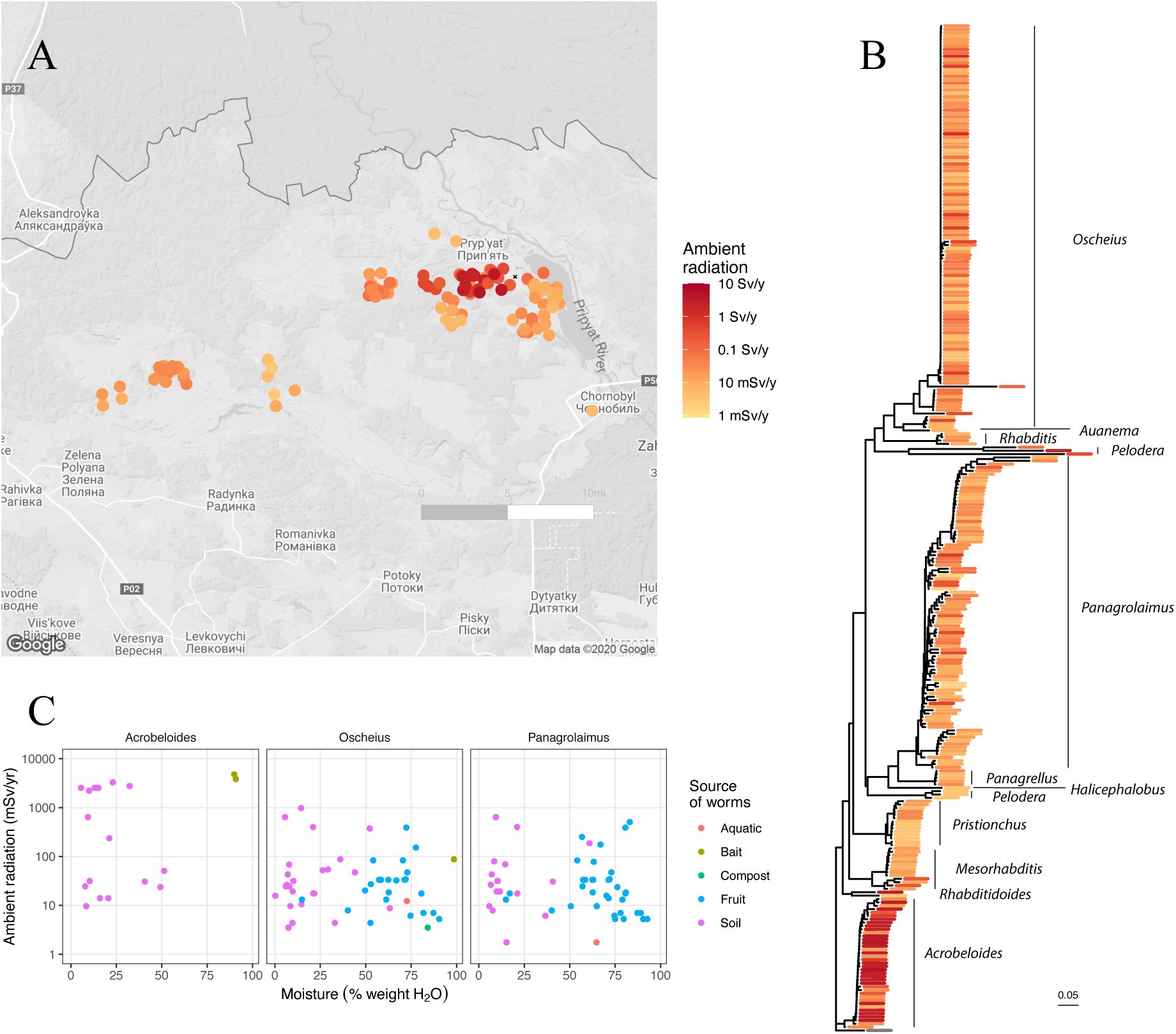
298 nematode lines collected, identified, and cryopreserved from the Chornobyl Exclusion Zone. (A) Sites from which substrates were collected. Jitter has been added to the positions to reduce overplotting. Colors represent ambient radiation at each site at the time of collection. (B) Neighbor-joining tree for a 518-bp segment 18S rDNA from 298 cryopreserved nematode strains. Branch lengths represent the proportion of sites that differ. (C) Environmental conditions at collection sites of the three most abundant taxa.

To identify each cultured isolate, we sequenced a ∼518 bp fragment of the 18S rDNA gene. All worms belong to Rhabditida, with representatives from *Acrobeloides*, *Mesorhabditis*, *Oscheius*, *Panagrellus*, *Panagrolaimus*, *Pristionchus*, *Pelodera*, and *Rhabditis* (Figure 1B). *Acrobeloides* were found more frequently in the drier and higher ambient-radiation environments, while *Panagrolaimus* and *Oscheius* were isolated from a broad range of radiation and moisture levels (Figure 1C). Collection information for all 298 cryopreserved samples is available in Supplementary File 1.

Of the three taxa we collected in the highest numbers, *Oscheius* was found in a broad range of environmental conditions (Figure 1B) and geographic locations (Supplementary Figure 3), and is the best characterized as a research model and most amenable to genetic analysis (*11, 24, 25*). For these reasons, we chose the *Oscheius* isolates to investigate further.

### *De novo* genome assembly of *O. tipulae* from Chornobyl reveal no genome rearrangements

Wondering whether the genomes of worms from Chornobyl carry signatures of living in a mutagenic environment, we selected 20 *Oscheius tipulae* for genome sequencing — 15 from the CEZ and 5 previously collected from other regions (Philippines, Germany, United States, Mauritius, and Australia). The 15 CEZ strains were selected (from a total of 120 *O. tipulae* isolates recovered from 53 unique sites) to represent a broad range of geographic locations and ambient radiation backgrounds, ranging from 3.5 to 978.3 mSv/y, across a distance of 50 kilometers. Additionally, we sequenced and assembled the genome of the JU75 strain of *O. sp. 3,* the sister species to *O. tipulae* (*26*), to use as an outgroup in subsequent analyses.

We performed long-read sequencing to facilitate *de novo* genome assembly, as well as short-read sequencing to polish the assembled genomes. For the 20 *de novo* genomes the average N50 was 9.14 Mb (median = 9.64 Mb, range = 5.85 Mb to 10.06 Mb), compared to the reference genome’s N50 of 10.82 Mb (*24*). *O. tipulae*, like *C. elegans*, is an androdioecious species and so are virtually homozygous, simplifying the genome assembly process.

Strains from Chornobyl are all genetically closer to each other than to the non-Chornobyl strains, with the exception of DF5103 from Berlin which clusters with the Chornobyl strains (Figure 2). Genetic distance between strains is correlated with geographic distance when compared across all strains, but not when compared among Chornobyl strains only (Mantel test with 999 replicates, observation = 0.930 and -0.021, p value = 0.001 and 0.478 for all samples and CEZ-only samples respectively, Supplementary Figure 4). Variants are distributed unevenly across the genome, with higher polymorphism in chromosome arms than centers (Supplementary Figure 5). Linkage disequilibrium drops off quickly (within 50 bases) but then plateaus around *r*^2^ = 0.4, reflecting a high rate of self-fertilization in this androdiecious species, while also suggesting that a that a low but nonnegligible rate of outcrossing (∼10^-3^-10^-6^/generation, (*27*)) does occur in the wild (Supplementary Figure 6).

**Figure 2:**
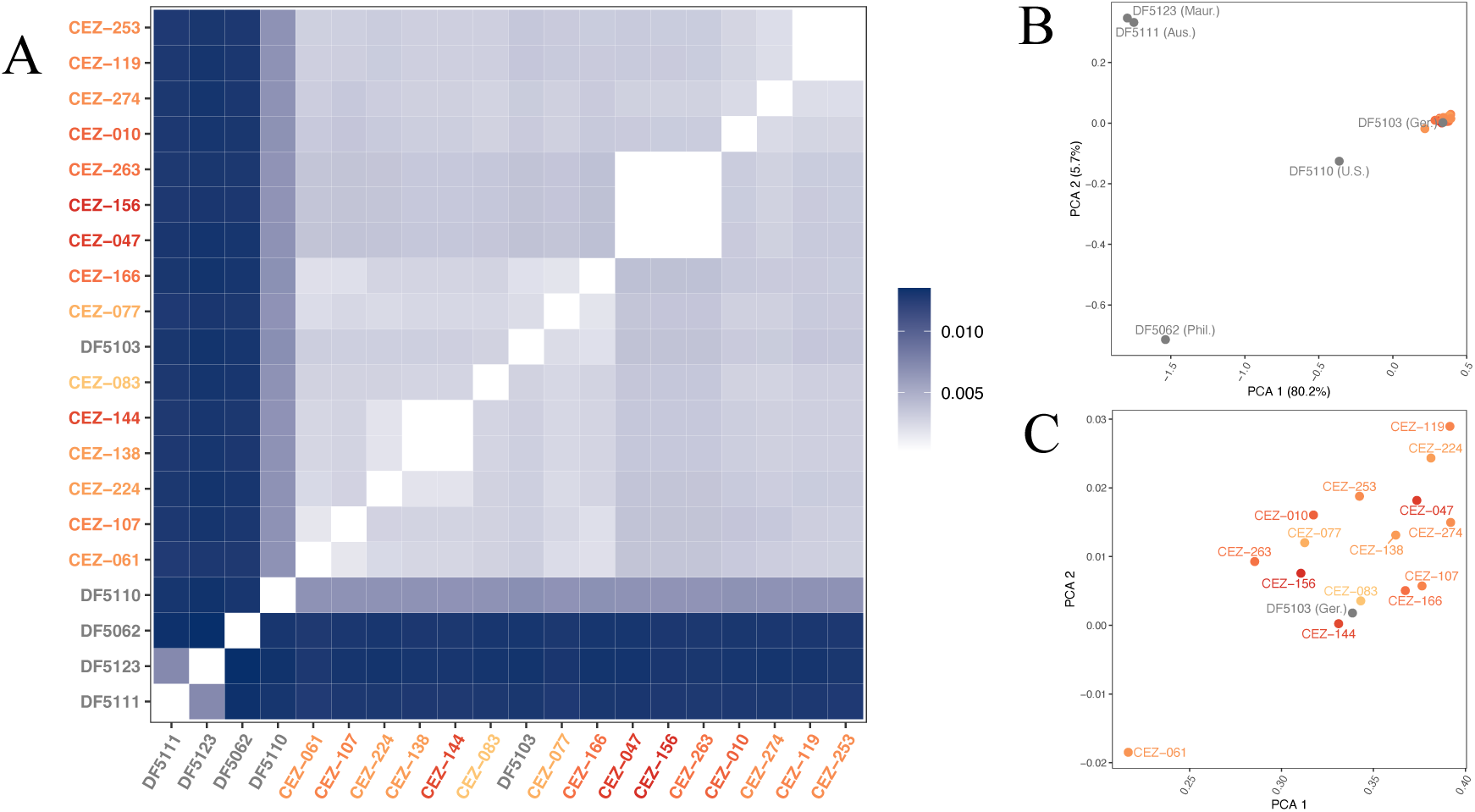
Population structure of 20 *O. tipulae* strains sequenced. (A) Genetic distances between each pair of samples. Values represent frequency of differences between the two strains. (B) PCA of 20 samples, and (C) inset of PCA in panel B to show detail of the Chornobyl and Berlin cluster.

Because ionizing radiation causes double-strand breaks in DNA, leading to rearrangements when resolved by non-homology-based mechanisms, we tested whether isolates from Chornobyl exhibited heritable chromosomal rearrangements. We generated synteny plots between the published reference genome (*24*) and each de novo genome (Figure 3). These synteny plots revealed no evidence of chromosome rearrangements compared to the reference genome, with the exception of a 2 Mb inversion on the left arm of chromosome V in non-Chornobyl strain DF5111 from Australia.

**Figure 3:**
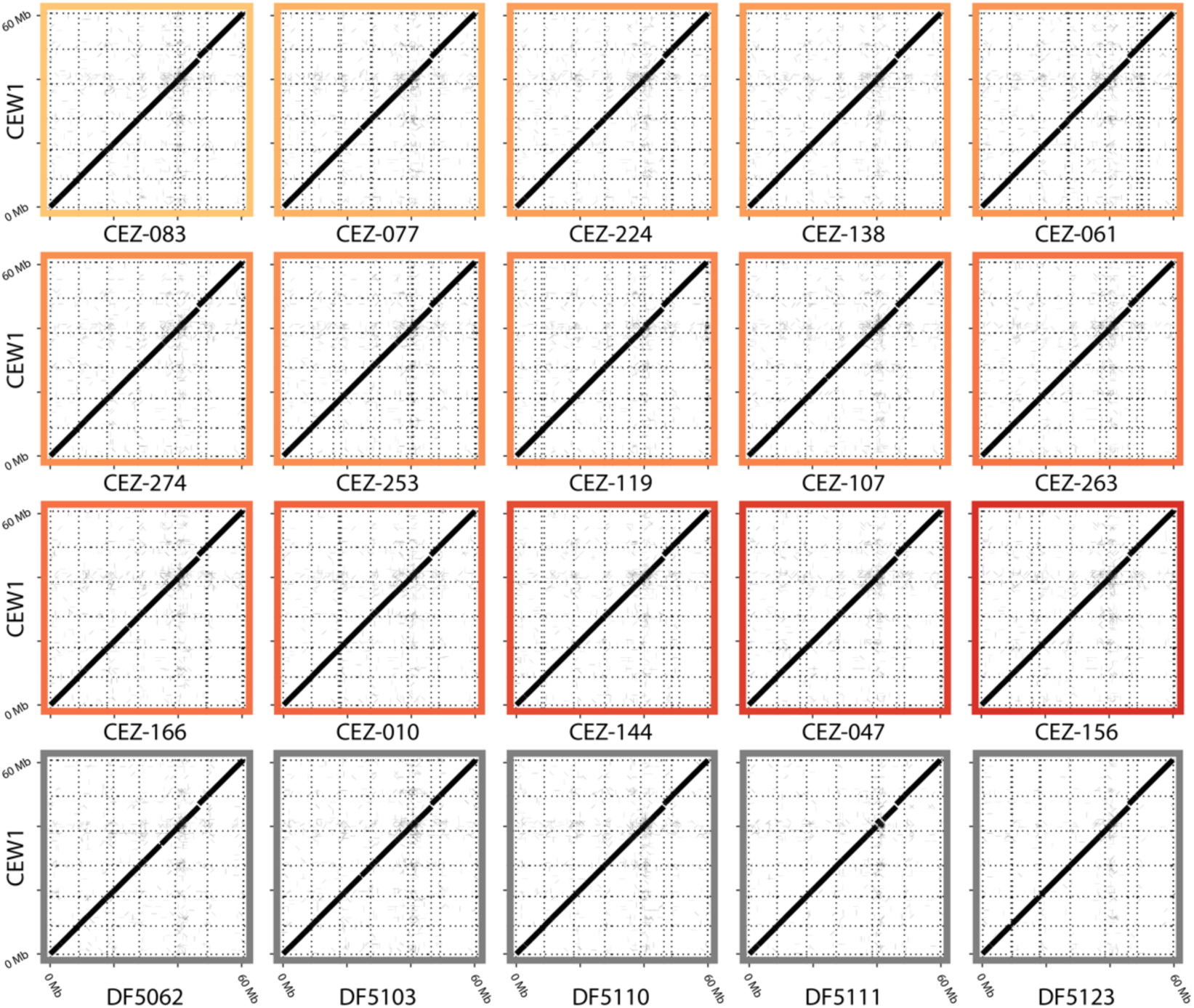
Synteny plots of each newly assembled *O. tipulae* wild isolate genome compared to the CEW1 reference genome. Border color reflects the ambient radiation at the time and place of collection, using the same color scale as in Figure 1. "DF" strains were collected previously, ambient radiations unknown, from Philippines (DF5062), Germany (DF5103), United States (DF5110), Australia (DF5111), and Mauritius (DF5123). Horizontal dotted lines delineate the six CEW1 chromosomes and vertical dotted lines delineate the contigs of each new genome. Stretches of alignment are plotted as lines that, due to the scale of the plot, are wider than they are long.

Differences in contig number between assemblies are explained by long-read coverage (Supplementary Figure 7), suggesting that the instances of low contiguity are artifacts of sequencing and not evidence of wide-spread chromosomal fission events. We observed a region of low synteny on the left arm of chromosome V, which could be explained by mobile sequences in all strains, rearrangements unique only to reference strain CEW1, or poor alignment in this region of the genome. Pairwise comparisons between every combination of genomes reveal similar patterns (example in Supplementary Figure 8), excluding the possibility that this is a CEW1-exclusive phenomenon. This region is coincident with the highest density of repeats in the genome (*24*). We inspected assemblies by eye to confirm that this high frequency of repeats was not leading to assembly errors, and we conclude that the low synteny in this region is likely real, and possibly caused by the high density of repeats (Supplementary Figure 9). The assembly gap on the right side of chromosome V in all new assemblies coincides with the long repetitive rDNA cluster.

### Recent mutations are comparable between worms from high-and low-radioactivity areas

Having found no evidence of large-scale chromosomal rearrangements, we next looked for signs that nematodes collected from higher radiation environments might have larger numbers of recently acquired mutations due to environmental exposure. To isolate recent mutations while accounting for variation in genealogy among strains and along the genome, we employed a tree-based method that uses an outgroup to assign recent mutations to branches in comparisons of two ingroup strains at a time (Figure 4A).

**Figure 4.**
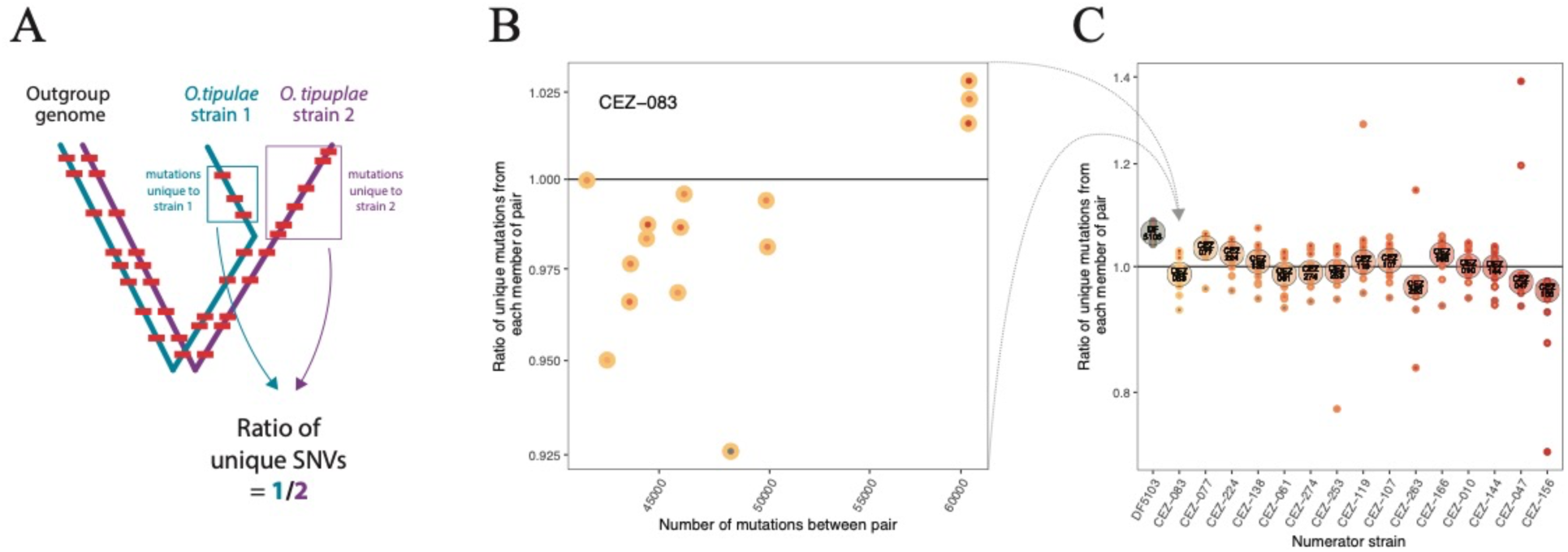
Relative mutation acquisition in *O. tipulae* wild isolates. (A) Schematic for calculating mutation acquisition of one isolate (strain 1) relative to another (strain 2). (B) Relative mutation acquisition for CEZ-083 when compared to each of the other 15 strains, for example. (C) Average relative mutation acquisition for each strain (p-value for glm testing association of average relative mutation acquisition to radioactivity = 0.187). Large circles are the median, and small circles are each pairwise comparison. Y-axis is plotted with a log transformation, tick labels report pre-logged values.

We aligned two *O. tipulae* genomes at a time to the genome of an outgroup, allowing us to count unique mutations in each of the two strains, accounting for variation in coalescence times along the genome. We define “relative mutation acquisition” as the ratio between these two unique mutation counts (Figure 4A). The goal of this approach is to disregard most differences each strain carries compared to the outgroup, and only count mutations acquired during recent divergence and whose ancestral allele can be determined from the outgroup, and then finally to report how much more one strain has acquired mutations compared to the other during the same period of unshared history.

Though *Oscheius sp. 3* is the closest known outgroup species, only 5-10% of the sites in each *O. tipulae* genome aligned to *O. sp. 3* genome. The *O. sp. 3* genome therefore is a mixture of conserved sites, likely under strong purifying selection, and sites that differ from *O. tipulae* by multiple substitutions; both factors reduce the utility of *O. sp. 3* as an outgroup. We therefore adopted an intraspecific outgroup approach, using *O. tipulae* DF5123 from Mauritius as our outgroup genome but limiting analysis to pairwise comparisons between European strains – a tight genetic cluster distant from DF5123 (Figure 2B) – and excluding genomic regions with genealogies inconsistent with an outgroup position for DF5123. With this intraspecific approach, 87-89% of the genome aligned to the *O. tipulae* reference and 78% of aligned variants fell within regions consistent with the desired genealogy.

In this way we generated 15 relative mutation acquisition values for each strain – one value for each comparison to the other strains (as an example, all pairwise comparisons using CEZ-083 in the numerator are shown in Figure 4B). We hypothesized that worms from higher radiation environments might have acquired more mutations in recent history than worms from lower radiation environments within the CEZ. Based on these 16 isolates (15 from Chornobyl and one from Berlin), we found no significant association between average relative mutation acquisition and radioactivity at the time of collection (p = 0.187 for ambient radioactivity by Geiger counter, p-value = 0.708 for substrate radioactivity by gamma spectrometer) for the nuclear genome (Figure 4C) or the mitochondrial genome (p=0.356 for ambient, p=0.818 for substrate). Comparing relative mutation acquisition to fold-change difference in ambient radiation levels between each pair using the Mantel test also revealed no association (p = 1), and subdividing data by mutation type also revealed no significant relationships (details in Supplementary Figure 10).

To test whether the Chornobyl strains differed systematically from non-Chornobyl *O. tipulae*, we repeated the analysis using all 20 sequenced strains with *O. sp. 3* as the outgroup. We saw no evidence of any significant difference in mutation acquisition between CEZ and non-CEZ strains (Supplementary Figure 11). Given the small and evolutionarily constrained fraction of the genome we were able to analyze, we consider this result inconclusive, and the earlier results from the German strain DF5103 (Fig 4C) provides the strongest evidence that Chornobyl strains lack signatures of elevated mutation accumulation.

The *O. tipulae* strains we sequenced include some that are very closely related to one another (0.025% divergence, compared to the range of 0.17-1.35% of all other samples). Of the approximately 2000 variants that differ between each of these pairs, two sets have a striking imbalance in how the variants are distributed: Among the 2,475 mutations differing between CEZ-047 and CEZ-156, they are disproportionately distributed at a ratio of 1.46:1 (binomial distribution p-value=1e-20), and the 2,881 sites that differ between CEZ-119 and CEZ-253 are distributed 1.28:1 (p-value=3e-11). The strong difference in recent mutation acquisition between these closely related strains is intriguing, but it is unclear if this effect is caused by demographic differences, differences in exposure history, or differences in each strain’s physiological response to mutagenesis. Differences in mutation spectrum between high and low mutation acquisition strains may suggest one mechanism over another, so we analyzed the similarities of each of these four mutation spectra to COSMIC signatures (*28*), but because there are only two pairs of highly similar strains, we were not able to determine statistical significance of these trends (Supplementary Figure 12). We were able to test our final hypothesis – whether strains have different physiological responses to mutagen exposure – in the lab using our cryopreserved cultures of each of these strains.

### *O. tipulae* strains exhibit heritable variation in their response to mutagen exposure, which is not explained by radiation levels at sites of collection

The result described above, that *O. tipulae* strains vary in their recent mutation acquisition but not based on their inferred radiation exposure, raises two questions that would be ideal to be able to verify or cross-reference via controlled experiments in the lab. First, why are Chornobyl *O. tipulae* no more mutated than non-Chornobyl *O. tipulae*? Is it because the level of radiation that they were exposed to was not challenging to the physiology typical of *O. tipulae*, or is it because the strains that thrive in Chornobyl are specifically more resilient to mutagens than other strains of the same species? Second, for the strains that do have fewer mutations in their recent history, as observed in the pairs described at the end of the last section, is it because they are more resistant to mutagenesis? Our cryopreserved cultures allow us to test these models directly. We tested isogenic descendants of the 20 natural strains of *O. tipulae* described above for their tolerance to mutagen exposure in the lab.

Motivated by previous experiments that showed reduced spermatogenesis to be the first dramatic fitness effects of ionizing radiation exposure (*29*), we developed a chronic mutagen tolerance assay that spans the entire lifecycle multiple times. We adapted the *C. elegans* Lifespan Machine method (*30*) to build an assay in which an isogenic population of worms is allowed to grow undisturbed for 2-3 generations with or without continuous exposure to a chemical mutagen, and population growth rates in both conditions are compared. Briefly, for each assay plate we added a consistent amount of bacterial food at high concentration and an isogenic population of synchronized L1 worms. We then imaged the plates continuously on flatbed scanners until the population size increased about 1 million-fold and the food was depleted from the plates. Because bacterial growth, and thereby worm food supply, might be affected by mutagen exposure as well, we fixed the bacteria with a quick, light paraformaldehyde treatment followed by washes. This preserved the bacteria cell shape - a requirement for some nematodes to successfully feed (*31, 32*). The number of hours it took each plate to exhaust its food is a proxy measurement for population growth rate broadly, and population growth rate is an umbrella fitness phenotype that we chose hoping to capture as many possible contributing fitness phenotypes as might be relevant (*e.g.*, brood size, time of onset of reproductive maturity, lifespan, body size, embryonic viability, etc.).

We used this multi-generational isogenic population growth rate assay to test 21 isolates’ sensitivities to three mutagens; cisplatin, ethyl methanesulfonate (EMS), and hydroxyurea (HU). Isolates included all 20 of the strains described above, as well as CEW1, the reference strain for *Oscheius tipulae* (*24*). Mutagens were chosen for their variety in DNA damage mechanisms, with EMS primarily causing bulky adduct formation (*33*), cisplatin causing inter- and intra-strand crosslinks (*34*), and HU causing replication fork stalling via depletion of nucleotides (*35*). For all three mutagens, strains differed in sensitivity (Figure 5 and Supplementary Figure 13).

**Figure 5.**
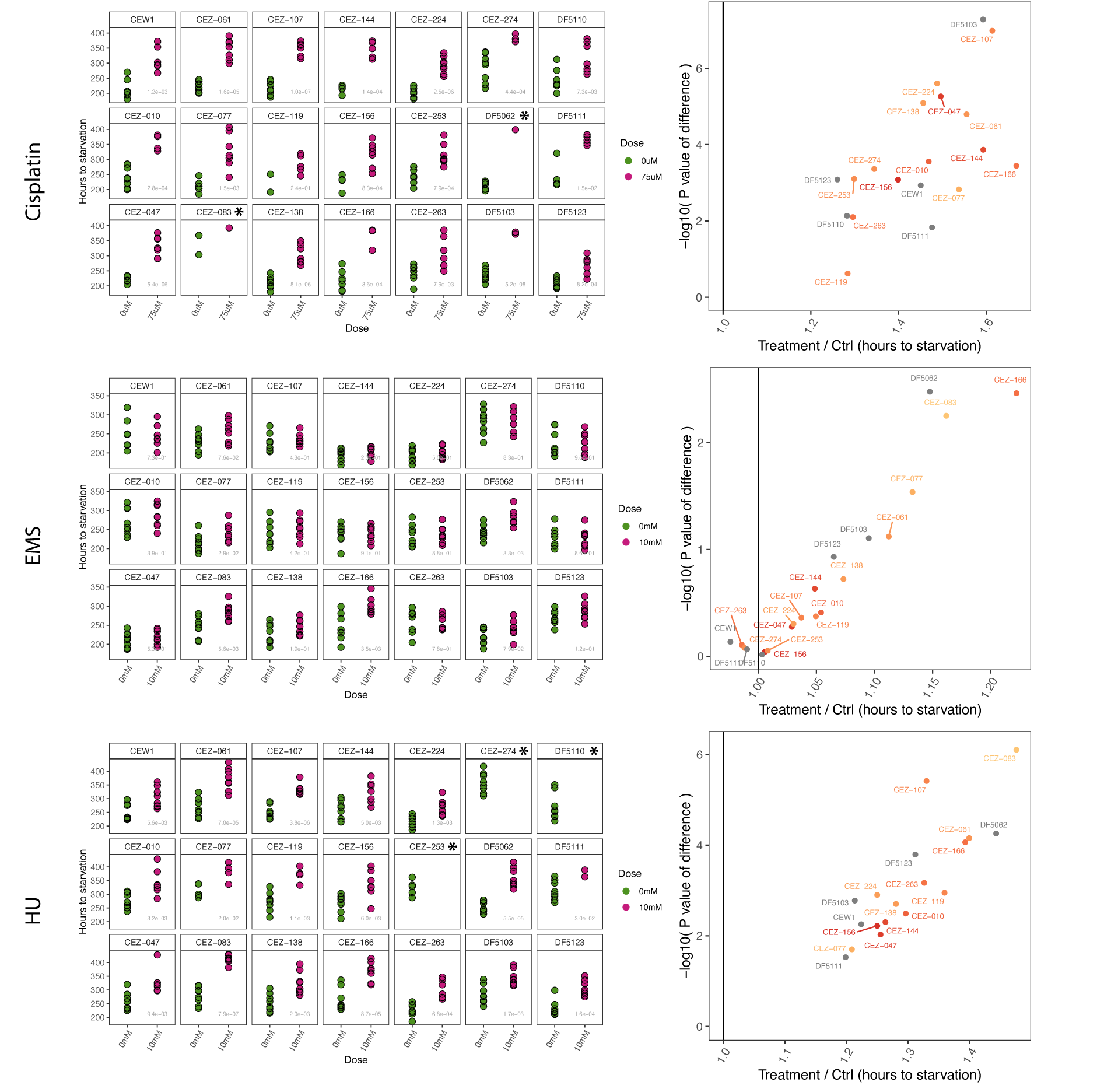
Mutagen tolerance assays of 21 strains of *O. tipulae* to three mutagens. Left: Hours of population growth until all food was consumed for cultures without mutagen (green) or with mutagen (pink). P-values for green vs. pink points shown in gray. Right: Ratio of average hours to starvation for mutagen-treated cultures compared to control cultures. Strains plotted further to the right are more sensitive, and strains plotted higher on the y-axis are more reproducibly sensitive. Asterisks in the left panels indicate strains for which 0 or 1 cultures consumed all the food on the plates with mutagen before the experiment was terminated. These strains do not appear in plots on the right.

We detected highly significant broad-sense heritability for sensitivity (*H*^2^*_sens_* = 0.37, 0.14, and 0.21 for cisplatin, EMS, and HU, p < 10^-10^ in all cases). However, we found no significant effects of Chornobyl versus non-Chornobyl origins (glm p-values for cisplatin = 0.596, EMS = 0.623, HU = 0.421), and no significant effects of ambient radiation dose within Chornobyl (glm p-values for cisplatin = 0.278, EMS = 0.144, HU = 0.155, Supplementary Figure 14). While mutagen tolerance cannot be broadly explained by the environment at the time and place of collection, there are nonetheless many interesting anecdotal observations from the intersection of these genomic and mutagen tolerance datasets (see Discussion), each of which can be pursued by genetic and cell biological experimentation in these strains.

Of the two sets of closely related pairs described at the end of the previous section, while the distribution of variants between them was unequal, none of these four strains were exceptionally tolerant or sensitive to the three mutagens tested (Figure 5). Therefore we were not able to attribute this different in mutation acquisition in the wild to broadly applicable differences in mutagen tolerance, and consider it more likely that this discrepancy is due to differences in the strains’ exposure history or demography.

Nematodes are highly robust animals, with some species reported as especially tolerant to harsh environmental conditions (*36–38*). To test whether *Oscheius tipulae* is a singularly tolerant species, we performed the same mutagen sensitivity assay at a range of concentrations in multiple strains of both *O. tipulae* and *C. elegans*, a species we did not find in the Chornobyl Exclusion Zone. *C. elegans* showed slightly higher tolerance to cisplatin, comparable tolerance to EMS, and slightly lower tolerance to hydroxurea (Supplementary Figure 15), leading us to conclude that *Oscheius tipulae* is not an exceptionally mutation-resistance species of rhabditid nematode.

## CONCLUSIONS/DISCUSSION

Microfauna from polluted or otherwise challenging environments provide valuable opportunities to investigate the nature of resilience and vulnerability within genetically diverse populations. These studies are often observational, as hypotheses and conclusions drawn from the field are not always testable within those same systems in the lab. In this study we characterized wild nematodes from sites of varying radioactivity within the Chornobyl Exclusion Zone, in the field and in the lab. We sequenced and analyzed the genomes of these isolates to investigate the genetic impacts of their recent environmental exposure, we cryopreserved cultures of each isolate, and we established a method for studying differences in tolerance to chronic exposure in this genetically manipulable species.

### Many taxa of nematode are abundant in the Chornobyl Exclusion Zone

We successfully isolated nematodes from sites of all levels of radioactivity throughout the CEZ. This may not be surprising – nematodes are robust animals with many life history-traits that can be advantageous in a hostile environment. Nevertheless, their reliable abundance in the wild and their experimental tractability position them well as a system that is portable between the field and the lab.

*Oschieus tipulae* emerged from our collections as the most abundant species from a broad range of environments. Also intriguing was the cephalobe *Acrobeloides*, which we collected in the highest frequency from the most radioactive sites. Parthenogenetic, and with a generation time of about a month compared to *O. tipulae*’s 5 days, *Acrobeloides* is less amenable to experimentation. But it has been previously described as a highly stress-tolerant animal (*36–38*) and may be interesting for future studies. Because the highest radiation sites for our collections were also distinctive in other ways (dry, exposed, and sandy), the prevalence of *Acrobeloides* there is not evidence that they are adapted to radiation.

### No evidence of Chornobyl-specific impacts on the genomes of *O. tipulae*

The genomes we assembled *de novo* from long- and short-read data revealed only a single large inversion, in a non-Chornobyl strain, suggesting that exposure-induced chromosomal rearrangements are not common in this population. Inspecting genomes for differential rates of smaller and more numerous mutations is not trivial, as counting private mutations in each strain would largely reflect the population structure of our samples, comparing each strain to a distant outgroup dilutes any effects of the last 35 years of exposure, and methods that use the full ancestral recombination graph require strong assumptions about unmeasured parameters.

We designed a strategy that compares the relative number of private mutations between each pair of strains, allowing us to identify which branch has acquired more mutations and which fewer in same periods of unshared history, for each pair. The results did not correlate with radiation levels at the collection site. Regardless, they do point us towards strains of interest, which are elaborated on in the next section.

### Mutagen tolerance varies heritably among strains, but Chornobyl nematodes do not show exceptional mutagen tolerance

One of the many appeals of the nematode as an experimental model is its versatility: It is a rare genetic system that is well-suited for studying individual cells, the entire animal, or populations of many millions of animals, all easily and affordably, within the lab. In this study we use a set of wild nematodes to bridge field work and lab experiments as well, by culturing Chornobyl strains and assaying their mutagen tolerance in a controlled setting. Comparing population growth over multiple generations in the presence or absence of three mutagens revealed heritably divergent responses to each mutagen. The fact that these responses vary independently of radiation levels at the site of collection suggests that the radiation these worms were exposed to likely did not surpass the limit of what standing variation in the *O. tipulae* population is equipped to survive. The possibility remains, though, that there are ionizing radiation-specific tolerance mechanisms selected for in Chornobyl that would be missed when testing for mutagen tolerance with the chemicals we used in this study.

By combining results from recent mutation acquisition in the field and mutagen tolerance experiments in the lab, several strains emerge as interesting for further study separately and in comparison. For each mutagen we now have relatively tolerant and relatively sensitive strains identified, between which we can compare the genetics and DNA integrity dynamics in response to mutagen exposure.

CEZ-263 is a strain with one of the highest tolerances to cisplatin and EMS, a moderate tolerance to hydroxyurea, and one of the lowest recent mutation acquisitions in the field compared to the other strains tested. These multiple data points support this strain’s candidacy as a broadly mutagen tolerant animal, and it will be interesting to compare it to a strain such as CEZ-166, which was most sensitive to cisplatin and EMS, moderately sensitive to hydroxyurea, and had one of the highest recent mutation acquisitions scores of those tested. These patterns are currently anecdotal, but they establish this species as a promising model for the genetics of DNA damage and repair.

## MATERIALS/METHODS

### Nematode collection and identification

We measured ambient radiation at each collection site by Geiger counter (Internal Medcom Inspector), averaging three measurements, each 1 cm above the ground and at least 30 cm from the others. To measure substrate radioactivity, we first estimated each sample’s moisture content by weighing the substrate, dehydrating with short pulses in a microwave or by leaving in an oven set to 38° C for 30 minutes at a time, and weighing again. Each sample was considered fully dehydrated once heat treatments stopped producing changes in weight. We then used a SAM940 portable gamma spectrometer (Berkeley Nucleonics Corp) with a 10 x 10 cm NaI detector to count Cs137 emissions. Gamma counts were performed inside a lead-brick enclosure to reduce background to under 10 counts per minute, and background was then subtracted from the sample spectrum. Counts were converted to corrected Becquerel as in (*39*).

All CEZ nematodes were collected using the Baermann funnel technique (*40*) conducted in Chornobyl within a day of sample collection. Taxonomic assignments are based on 18S sequence similarity to nematodes in the NCBI nucleotide database. A region of 18S was amplified and sequenced using RHAB1350F (5’-TACAATGGAAGGCAGCAGGC) and RHAB1868R (5’-CCTCTGACTTTCGTTCTTGATTAA) primers (*41*). The gene tree in figure 1B was generated by aligning 18S sequences in R (*42*) using msa() from the *msa* package (*43*) with default settings, then using *ape* (*44*) to calculate distances (dist.dna(model = "raw", pairwise.deletion = T)), estimate a tree (njs()), root the tree with the *Plectus minimus* sequence (root()), and plot (plot.phylo()).

### Long-read DNA sequencing

Worms were grown on 10 10-cm plates at 20° on NGMA, washed for one hour in M9, then quickly washed with sterile distilled water, pelleted, and resuspended in 5x volume lysis buffer (200 mM NaCl, 100 mM Tris-HCl pH 8.5, 50 mM EDTA pH 8.0, 0.5% SDS, 0.1 mg/mL proteinase K added fresh). Worms were lysed at 65°C for 3-12 hours, then incubated at 95° for 20 minutes. RNase A was added at 0.1 mg/mL and incubated for one hour at 37°. DNA was isolated by mixing the sample, phenol, chloroform, and isoamyl alcohol in 50:25:24:1 proportions and shaking for 20 seconds, then spinning down in phase-lock gel tubes for one minute at 4000 rcf, followed by 5 minutes at 16000 rcf, and keeping the top aqueous phase. Ethanol precipitation was performed by adding 0.1x volume 3M NaAc and inverting, then adding 1 mL (or >2x total volume) 100% ethanol, mixing, incubating for 20 minutes at 4°, centrifuging for 30 minutes at 4° at 16000 rcf, removing the supernatant, washing the pellet twice with 70% EtOH, then drying the pellet and resuspending the DNA overnight in 100 µL water. Size selection was performed with the Short Read Eliminator Kit (Circulomics #SS-100-101-01) according to the manufacturer’s instructions. Purified DNA was prepared and sequenced using the Oxford Nanopore Technology MinION device (Ligation Sequencing Kit SQK-LSK110, Spot on flow cell Mk1 FLO-MIN106D, and Native Barcoding Expansion EXP-NBD104) according to the manufacturer’s instructions.

### Short-read DNA sequencing

DNA was extracted by salting out (*45*). Worms were rinsed in M9 for one hour then pelleted and frozen overnight. The pellets were thawed and incubated in 275 µL TNE with 7.5 µL 20% SDS and 15 µL 20 mg/mL proteinase K at 55° overnight. The next day samples were vortexed, another 15 µL proteinase K added, and incubated at 55° for another two hours. After 6 µL 100mg/mL RNase A was added, samples were vortexed and let rest for two minutes. 85 µL 5M NaCl was added and samples were shaken vigorously. Samples were spun at 14000 rpm for 10 minutes and the supernatant was moved to a fresh tube. 350 µL ice cold EtOH was added, and tubes were inverted and then spun at 14000 rpm for 3 minutes. Supernatant was removed and the pellet dried and resuspended in 20-100 µL TE at 37° for a few hours. DNA was passed through a Genomic DNA Clean & Concentrator kit (Zymo D4065) and prepared for sequencing using the NEBNext FS DNA Library Prep Kit (E7805) according to the manufacturer’s instructions. Libraries were sequenced on a NextSeq 500 for 2x150 cycles at the NYU GenCore Facility.

### Genome assembly

Bases were called and barcoded from MinION data using Oxford Nanopore’s guppy/4.4.0_gpu (guppy_basecaller with a minimum qscore of 7, guppy_barcoder). Contigs were assembled with Flye v2.8.1 (flye --nano-raw --genome-size 60m --iterations 1 -- meta) (*46*), then cleaned with Oxford Nanopore’s Medaka (medaka_consensus -m r941_in_high_g360) and Racon v 1.4.19 (*47*) using default settings. Illumina reads were aligned to post-Racon assembly using minimap2/2.17 (minimap2 -ax sr) (*48*), and alignments were used to polish assemblies with pilon-1.24 (pilon --geno) (*49*). Polished contigs were inspected using blobtools v1.1 (map2cov, create, view, and plot) (*50*), and contigs that were likely contaminants due to their copy number and/or GC content were excluded manually from the final assembly. In three cases chromosomes were assembled in what appeared to be chromosomal fusions, but inspection of long-read alignments revealed zero reads that spanned non-telomeric regions of both sides of the fusion, so this was inferred to be an assembly error and contigs were manually split.

### Genetic distance, PCA, and Mantel test

We subsampled Illumina reads for each isolate’s genome so that all libraries had similar coverage (∼29x). These subsetted Illumina reads for each sample were aligned to CEW1 reference genome using minimap2 v2.17 and samtools v1.11 (*48, 51*) (minimap2 -ax sr -R [manually added readgroup], samtools view -S -b, samtools sort, samtools index). Alignments were genotyped and converted to vcfs using GATK v4.3.0.0 (HaplotypeCaller -ax sr to generate GVCFs, GenomicsDBImport to combine all samples into one database, GenotypeGVCFs -all-sites to produce a vcf) (*52*). Genetic distances were generated by selecting two strains at a time, filtering them for only sites in which both samples had a read depth >5, and counting the fraction of sites that differed between the pair using bcftools v1.14 (*53*) (bcftools view -s strain1,strain2 -m 2 -M 2 -i ’MIN(FMT/DP)>5’). PCA was performed using sites for which all samples had a read depth >5. Mantel tests were performed using the *adegenet* (*54*) R package command mantel.randtest() on genetic distance calculated above and haversine geographic distance calculated with the *geosphere* (*55*) package (distHaversine()).

### Synteny plots

Synteny plots were generated by aligning assembled genomes to the CEW1 reference genome or to each other with minimap2 v2.17 (minimap2 -x asm20) (*48*), and were plotted in R using the package pafr (*56*).

### Relative mutation acquisition

We subsampled Illumina reads for each isolate’s genome so that all libraries had similar coverage (∼29x). We then aligned reads to the JU75 genome (outgroup *O. sp 3*) or the CEW1 genome (*O. tipulae* reference) using minimap2 v2.17. We called SNPs and indels with GATK v4.1.9.0 (HaplotypeCaller -ax sr, GenomicsDB, GenotypeGVCFs).

To avoid including genomic regions for which DF5123 might not be a suitable outgroup, we broke the full vcf table into 1000-variant pieces and generated a distance matrix for each piece. If a set of 1000 variants generated any DF5123 distances that were less than any non-DF5123 distances, we excluded all 1000 variants from this analysis. By this method we removed 22% of variants.

For each possible pair of samples, we made a vcf of those two strains plus DF5123 (the intraspecies outgroup) mapped to CEW1 (the annotated reference strain for *O. tipulae*). We filtered for only sites with two alleles and a minimum read depth of 5 (bcftools v1.14, bcftools -s strain1,strain2 -m 2 -M 2 -i ’MIN(FMT/DP)>5’). From this pruned and filtered vcf we counted the ratio of mutations unique to each sample for each mutation type, added 50 count to each to buffer against inflated effects of small numbers, and then divided the counts from strain 1 by the counts from strain 2 to generate a relative mutation acquisition value.

The Mantel test was performed using the *adegenet* (*54*) R package command mantel.randtest(nrepet=999) on ambient radiation fold change and relative mutation acquisition value for each comparison in both directions.

### Fixing food for mutagen experiments

Bacterial cultures of *E. coli* NA22 were grown in Terrific Broth overnight at 37° and *Ochrobactrum vermis* MYb71 (*57*) were grown in Terrific Broth at 25° for two nights, all in 1-liter beveled flasks. Cultures were spun down in a Sorvall floor centrifuge in 1-liter bottles at 5000 rpm for 10 minutes. Pellet weights were determined by weighing empty Sorvall bottles before adding medium and weighing them again after spinning down and removing supernatant. Pellets were resuspended in sterile water 3x the mass of the pellet. Paraformaldehyde was added to reach a final concentration of 5% and incubated shaking at 37°C for 1 hour. The solution was spun down again in a Sorvall centrifuge, this time in 50 mL tubes, and once again resuspended in 3x the mass of the pellet using sterile water. These two stocks of fixed bacteria were mixed together in a 1:1 ratio by volume to produce the final food stock for mutagen sensitivity assays.

### Mutagen sensitivity assay

Worms were maintained at 20° on NGMA plates seeded with OP50 *E. coli*. Gravid adults were bleached on Day 1 and embryos were plated without food to induce L1 arrest. On day 2, synchronized larvae were counted by averaging the number of larvae and late-stage embryos per 1 µL drop for 3 drops, and roughly 50 worms were added to each 3.5 cm NGMA plate with 10 µg/mL nystatin and 50 µg/mL streptomycin. Eight plates of the negative control condition and eight plates of the experimental treatment condition were prepared for each strain. Stocks of fixed NA22 and MYb71 bacteria (see above) were mixed 1:1 by volume, and mutagen solutions (resulting in final concentrations 75 µM cisplatin, 10 mM EMS, or 10 mM HU) or water (for controls) were added to aliquots, 200 µL of which were then distributed onto each plate. Concentrations of mutagens were calculated based on a final volume of the entire 35mm NGMA plate (3.31 mL), assuming that the mutagen is diffused through the entire plate for the majority of the experiment. Plates were randomized and arranged on acrylic racks on Epson V750, V800, and V850 flatbed scanners in a 20° incubator. Acrylic racks held 32 plates in place on each scanner, with 9 mm ventilation below and 4.5 mm ventilation above plates. Scanners collected 1200 dpi images once per hour continuously until all plates had starved or begun to dry out, usually 10 days for *C. elegans* and 20 days for *O. tipulae*. The standard deviation of pixel intensities from each image was calculated by ImageJ and values were analyzed in R to determine the exact time of food depletion, which was identified by with the abrupt end of exponential population growth and sudden drop in standard deviation of pixel intensity.

### Statistics for mutagen sensitivity assay

We estimated broad-sense heritability of toxin-sensitivity using a linear mixed-effect model (*58*). The phenotype is hours to starvation, and the data (after censoring plates that became contaminated or never reached starvation) include 18-21 strains measured on 1-8 plates per strain per treatment, randomized across eleven flatbed scanners, all assayed in a single experimental batch for each toxicant. Fixed effects are the hours-to-starvation intercept and the effect of treatment (toxin vs control). Random effects are worm strain (random intercepts), treatment effect within strain (random slopes), experimental scanner (random intercepts), and residual microenvironmental variation. The plate-level broad-sense heritability of sensitivity is the fraction of random-effect variance attributed to treatment within strain (i.e., variance explained by the random slopes). Note that among-strain variation in baseline population growth rates (i.e, variance explained by random strain intercepts) accounts for an additional 0.27, 0.37, 0.26 of the random-effect variance in the cisplatin, EMS, and HU experiments respectively.

We tested significance of the heritability of sensitivity by comparing models with and without random slopes, using likelihood ratio tests and the Chi-square distribution to estimate p-values.

## SUPPLEMENTARY FILES

1. QG numbers and collection conditions for 298 cryopreserved samples - <u>link</u>
2. VCFs - <u>link</u>
3. Mutagen sensitivity data - <u>link</u>

## DATA AVAILABILITY

Sequence data has been uploaded to NCBI (fastqs to SRA and genomes to WGS) under project accession # PRJNA974388.

## CODE AVAILABILITY

All R code, and the necessary supplementary data files, will be available at https://github.com/tintori/chornobyl_genomes. Currently the files are available at https://www.dropbox.com/sh/61wsg37cn15be8u/AABDV8OZ0bxm17wJookZMZ3ea?dl=0.

## AUTHOR CONTRIBUTIONS

Experimental design: ST, MR

Field work: ST, IC, TM, MR

Culture and cryopreservation: ST, PO, MR

Tolerance experiments: DC, ST

Analyses: ST, MR

Manuscript prep: ST with feedback from MR

## ACKNOWLEDGEMENTS

This work was supported by grants ES031364, ES029930, GM141906, Damon Runyon Cancer Research Fellowship DRG-2371-19 (to ST), a grant from the Zegar Foundation to the NYU GenCore Facility, and from the Samuel Freeman Charitable Trust (to TM). We thank Maxim Ivanenko for sharing his invaluable expertise in the field, Jonny Turnbull for help collecting nematodes, and Karin Kiontke for help interpreting 18S sequences. Thanks to Erik Andersen, Ryan Baugh, members of the Andersen, Baugh, and Rockman labs, Sevinç Ercan, and Molly Przeworski for helpful conversations. Thanks to NYU Rhabditid Collection for all DF *Oscheius* strains, to Marie-Anne Félix for the JU75 strain, and to the *Caenorhabditis* Genetics Center, funded by the NIH Office of Research, Infrastructure Programs (P40 OD010440), for bacterial strains NA22 and MYb71.

## SUPPLEMENTARY FIGURES

**Supplementary Figure 1.**
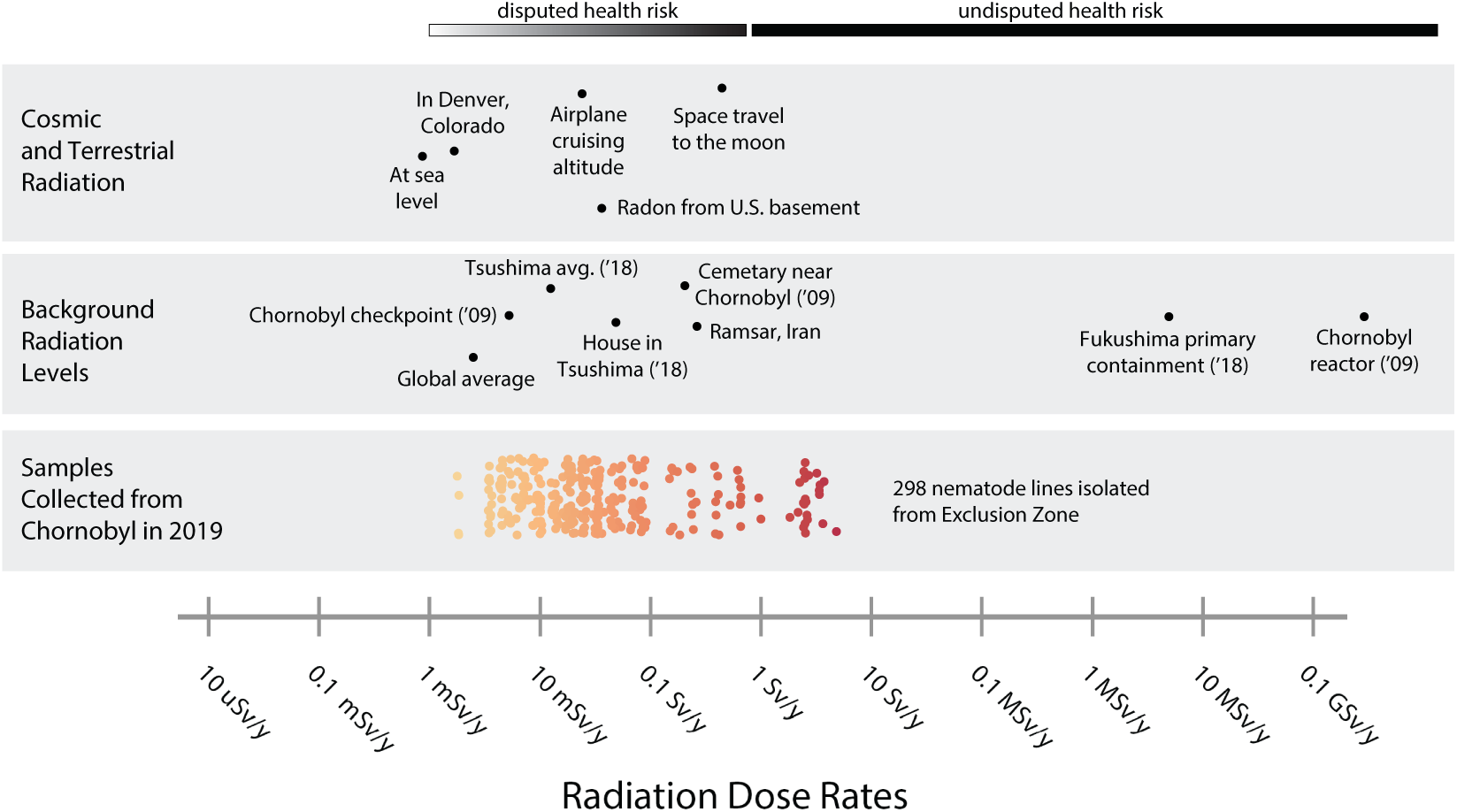
Ambient radiation levels of nematode samples collected from the Chornobyl Exclusion Zone, compared to reference points. Data aggregated from Supplementary References (*59–67*).

**Supplementary Figure 2.**
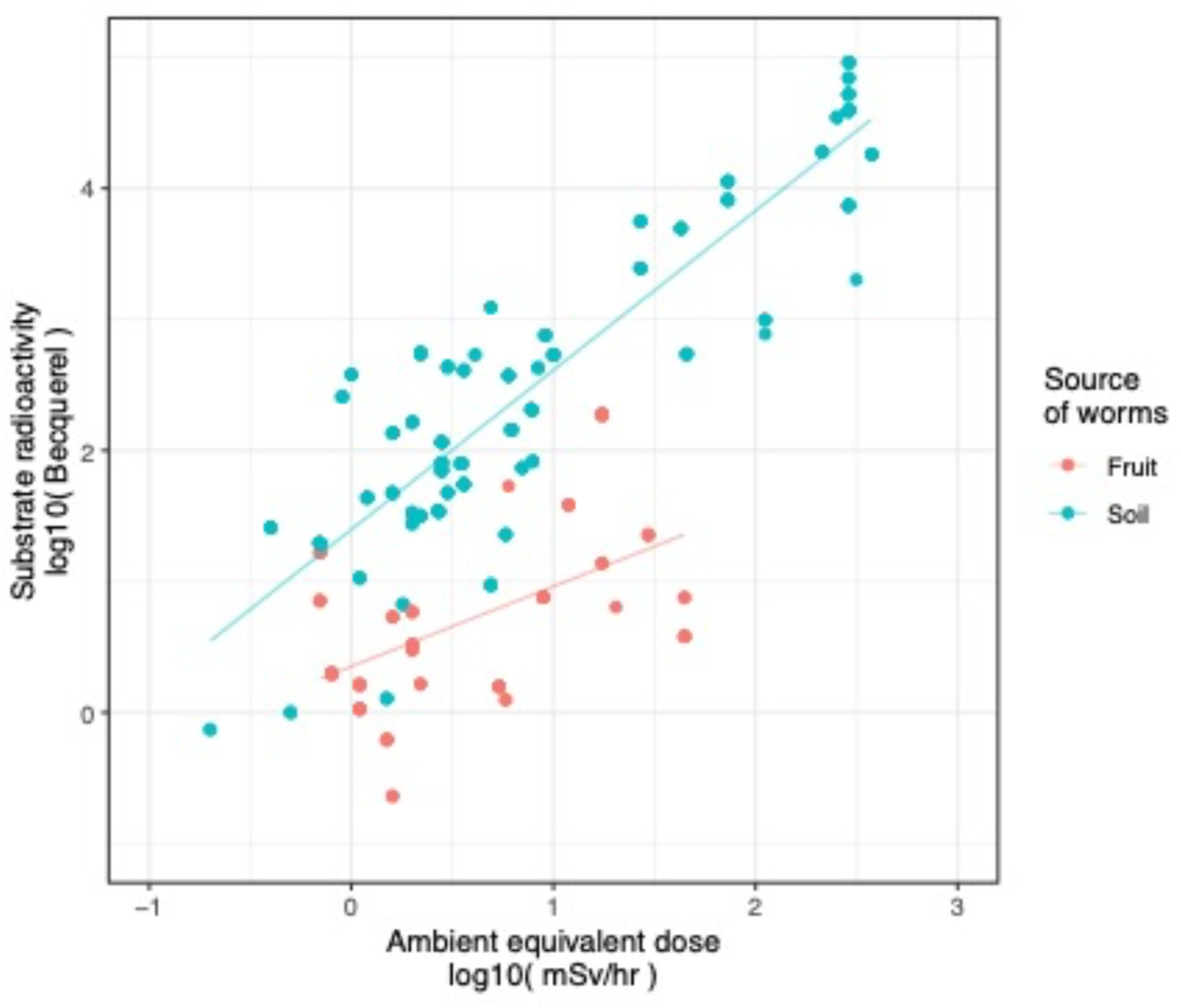
Substrate radioactivity increases as environmental radiation increases. Fruit samples (red) have similar substrate and ambient radiation levels whereas soil samples (blue) tend to be more radioactive than the ambient radiation in the environment from which they were collected. Ambient radiation values were generated by averaging three Geiger counter measurements 1 cm from the ground, >30 cm apart from each other. Substrate radiation values generated by gamma spectrometer.

**Supplementary Figure 3.**
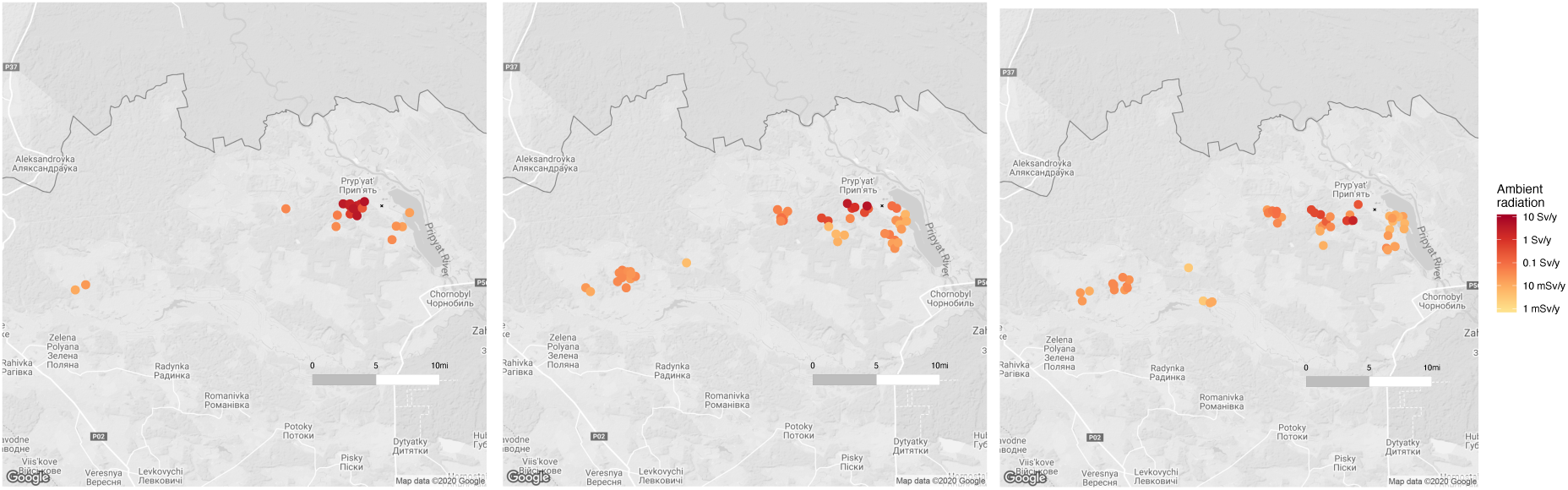
Locations and ambient radiation levels of collection sites from which *Acrobeloides* (left), *Oscheius* (center), and *Panagrolaimus* (right) were collected.

**Supplementary Figure 4.**
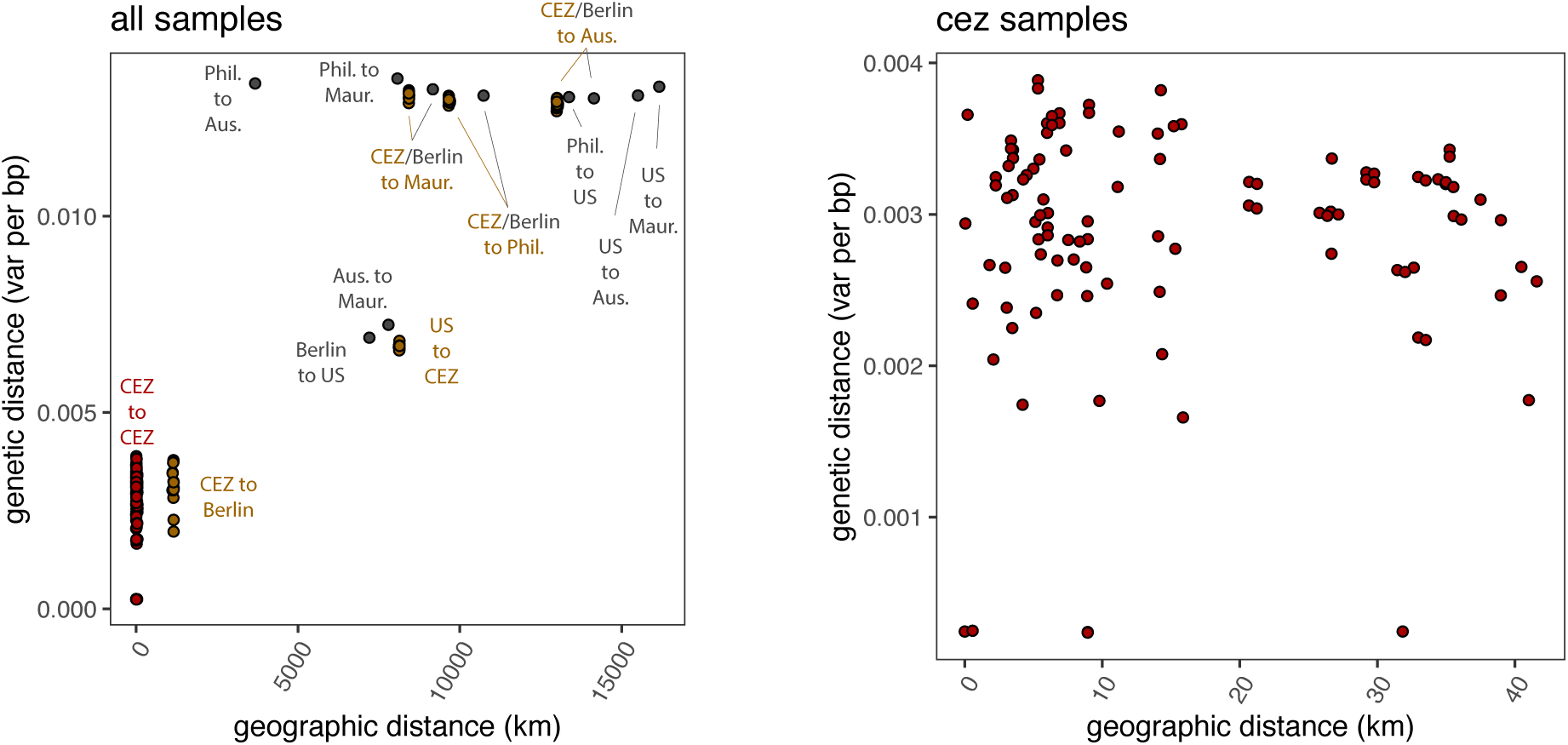
Genetic distance correlates with geographic distance when including all possible samples (left) but not when looking within Chornobyl strains exclusively (right). Each dot represents a pairwise comparison between two strains. Red: Chornobyl to Chornobyl comparisons, gray: non-Chornobyl to non-Chornobyl, brown: Chornobyl to non-Chornobyl. Plot on the right shows all the red points from the bottom left corner of the plot on the left.

**Supplementary Figure 5.**
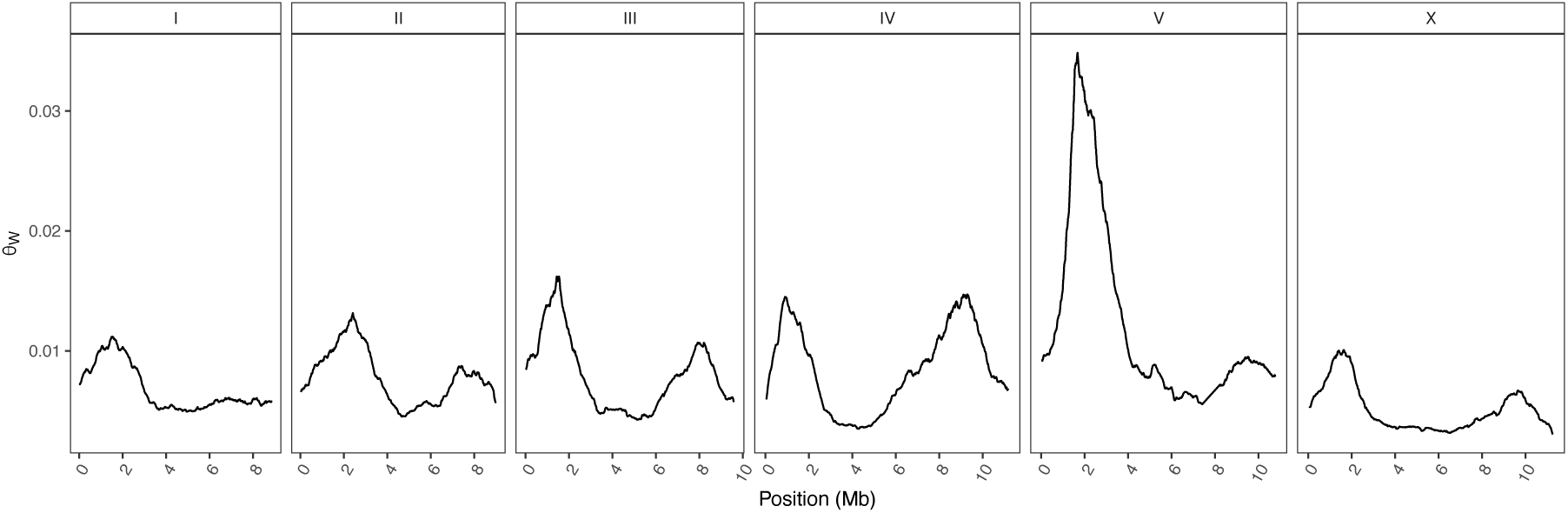
Variant distribution across the *O. tipulae* genome. Watterson’s theta calculated for a 1 Mb sliding window centered every 10 kb. Theta was calculated in R by dividing the number of varying sites by the number of sites at which all 20 samples had a read depth greater than 5, for 1 Mb windows centered every 1kb. This ratio was divided by the sum of 1/i for i=(1..19).

**Supplementary Figure 6.**
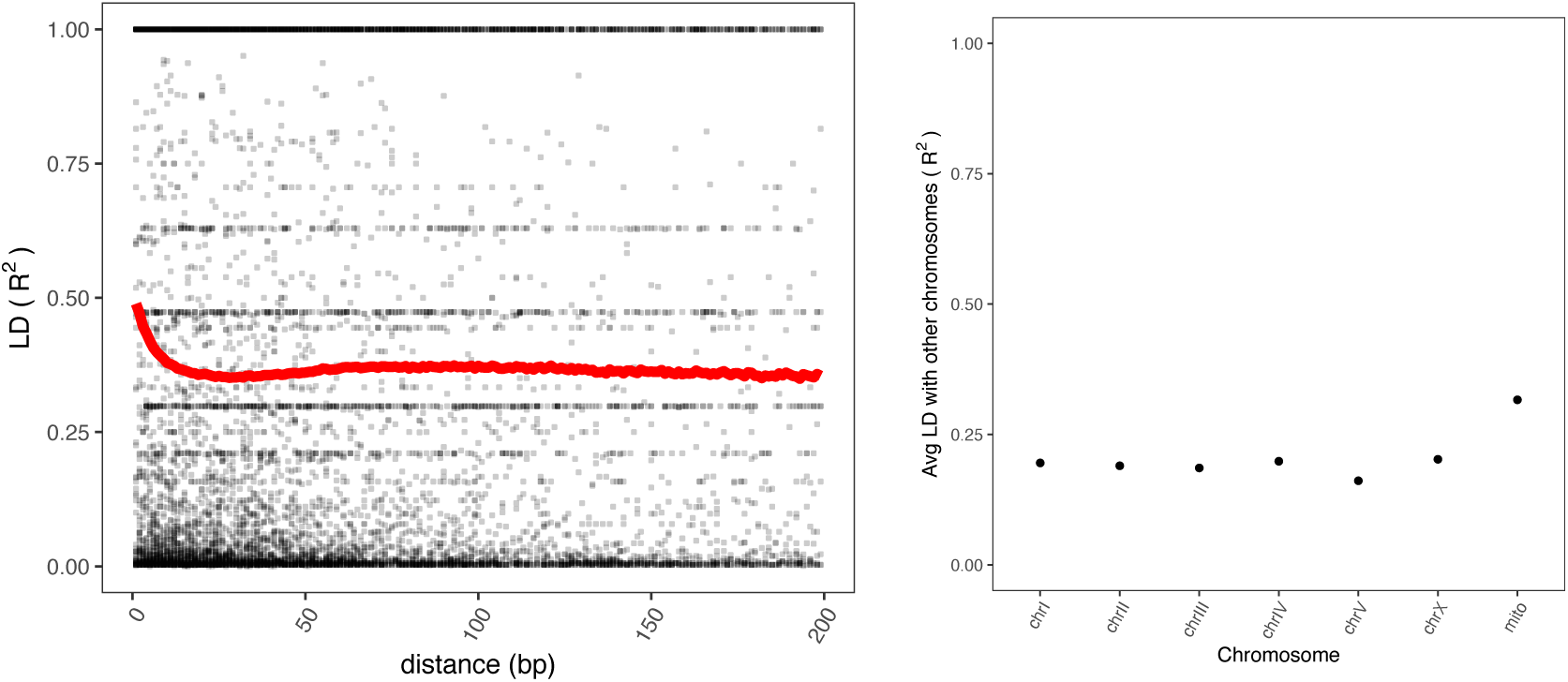
Linkage disequilibrium within nuclear chromosomes (left) and between chromosomes (right) of *O. tipulae.* R^2^ was calculated using plink (*68*) v1.90b6.21 with --ld-window-r2 0 for both panels, and the VCF thinned to 0.5% for the right panel (between chromosome analyses).

**Supplementary Figure 7:**
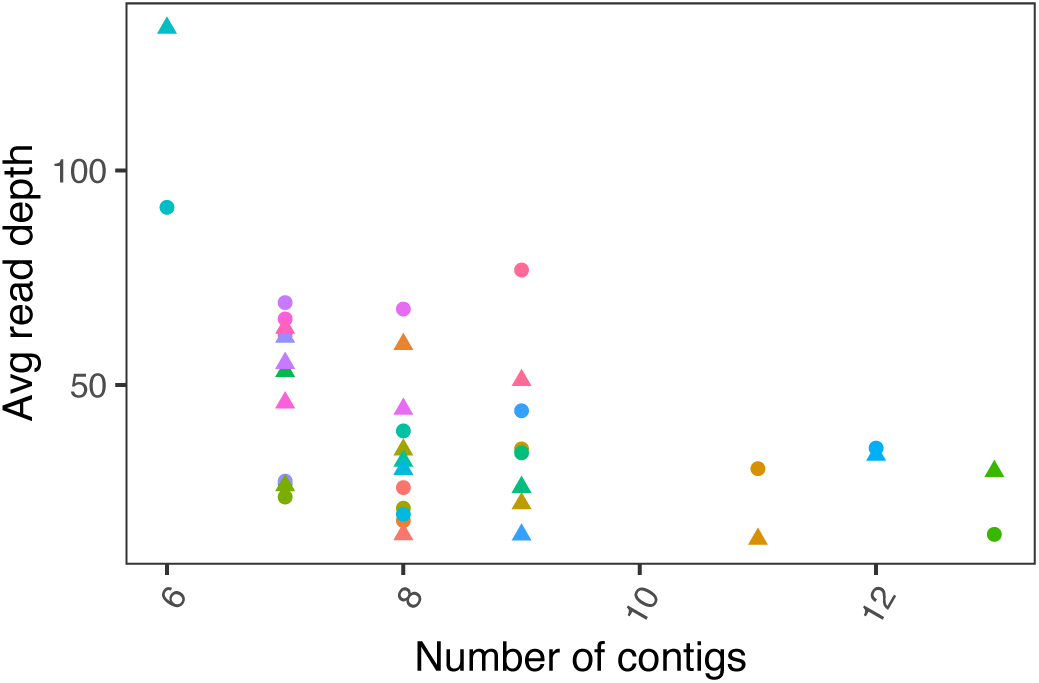
Assembly contiguity can be partially explained by coverage of long-read (Oxford Nanopore Technology MinION) sequences (*p* = 0.019). Higher-contiguity assemblies (towards the left, with fewer and larger contigs) correlate with deeper Oxford Nanopore sequencing. Only contigs larger than 200K were counted, disregarding small pieces that did not assemble well.

**Supplementary Figure 8:**
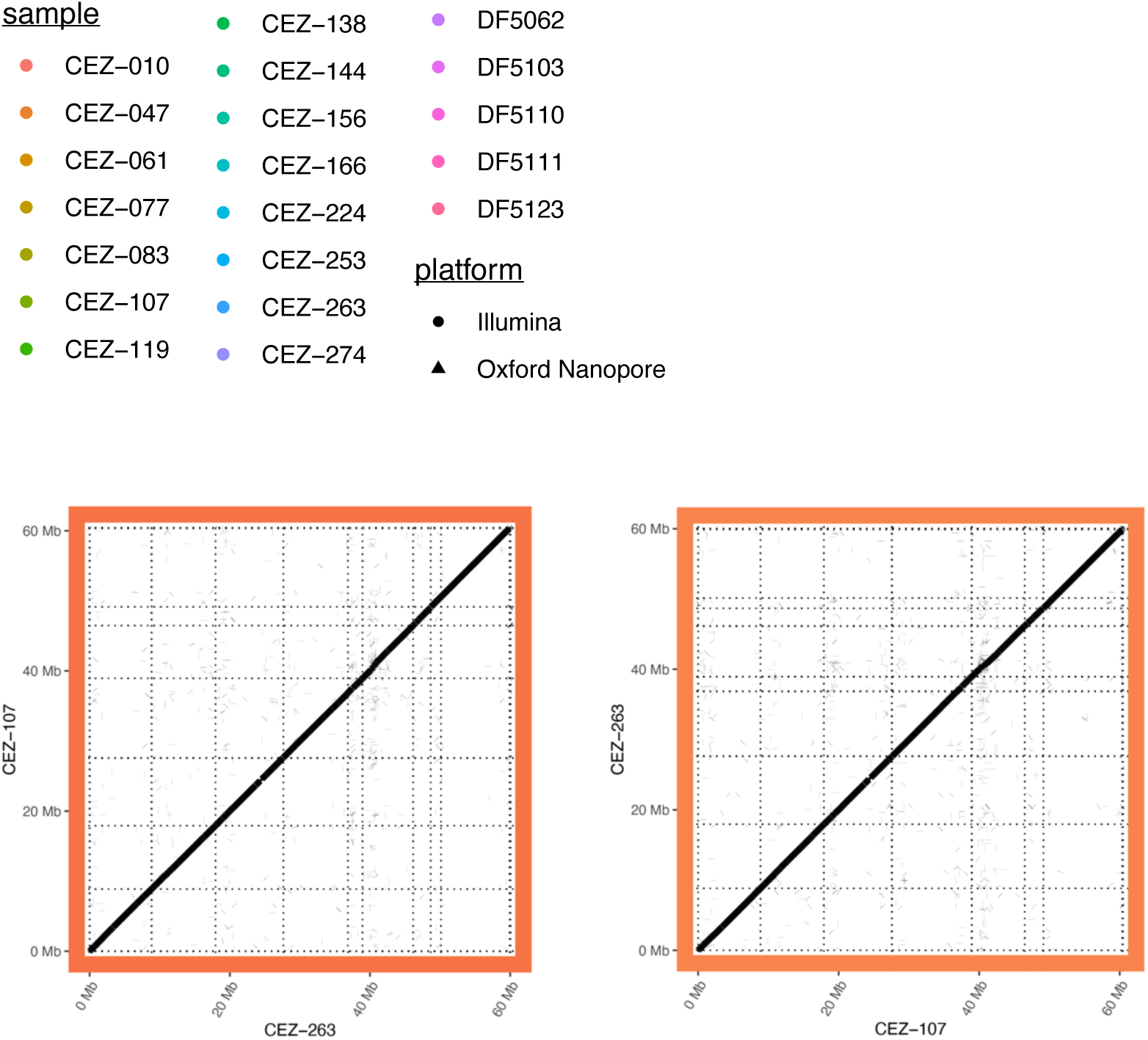
Synteny between CEZ-107 and CEZ-263. The low synteny region on chr V persists between *de novo* genomes, not just when compared to CEW1 reference as in figure 3, implying that the chr V smear is not showing CEW1-specific rearrangements. (Left) CEZ-107 or (right) CEZ-263 as reference genome. Strains were randomly selected from all non-CEW1 *O. tipulae* genomes.

**Supplementary Figure 9.**
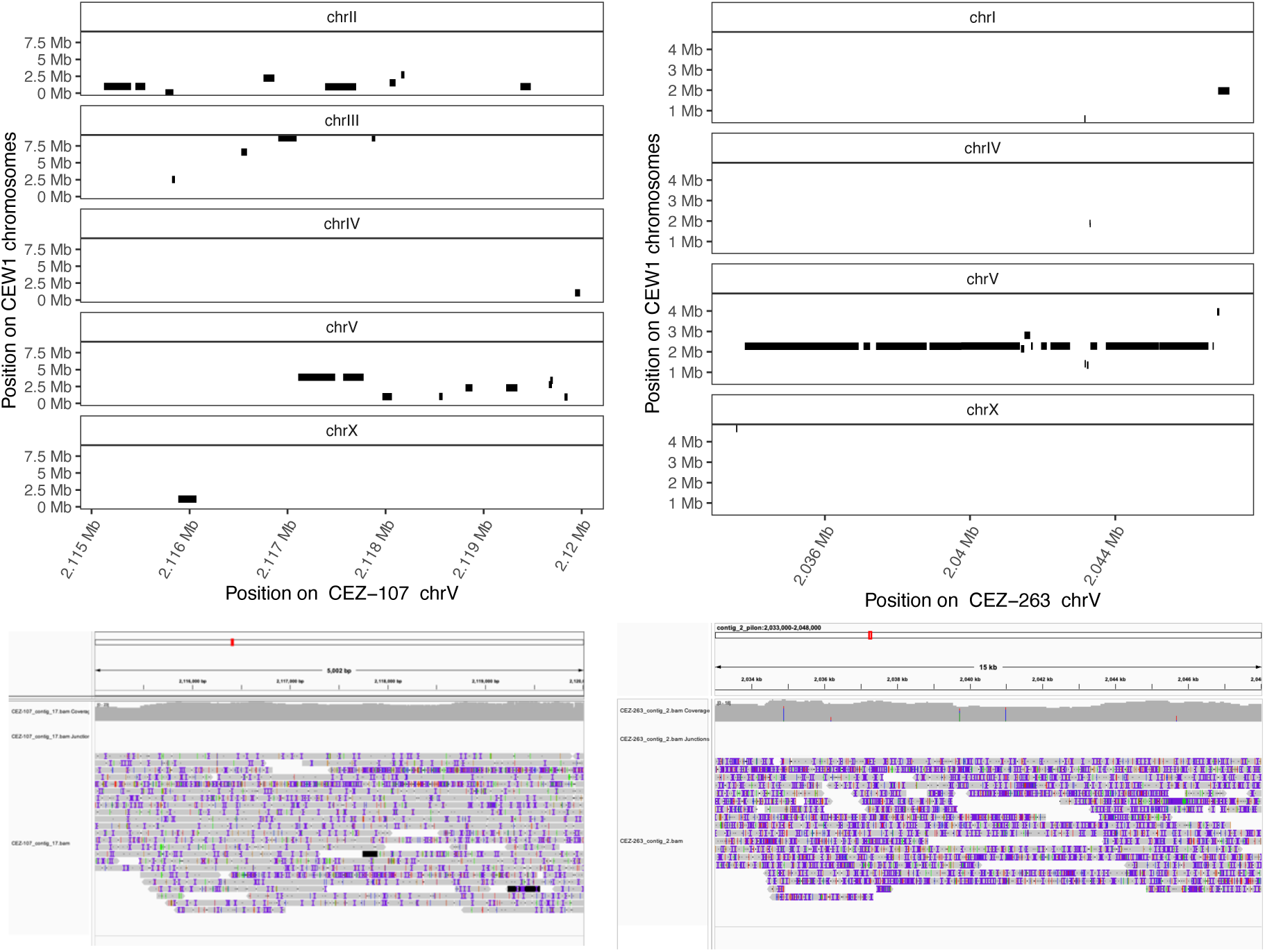
Visual inspection of assembly over the low synteny region on chr V. Top: Synteny plots show low-synteny aligned fragments. Within a 5 kb interval of CEZ-107 chr V (left), small fragments map to many different regions of the CEW1 genome. Within a different 15 kb interval of CEZ-263 chr V (right), synteny with CEW1 chromosome V is largely conserved. Bottom: Long-read pileups of these same regions show that the long reads continuously span this low-synteny region. This confirms that poor synteny on chr V is not due to low coverage or abrupt misassemblies in genomes. Each gray bar represents a long read, with polymorphisms between the two genomes represented in color.

**Supplementary Figure 10:**
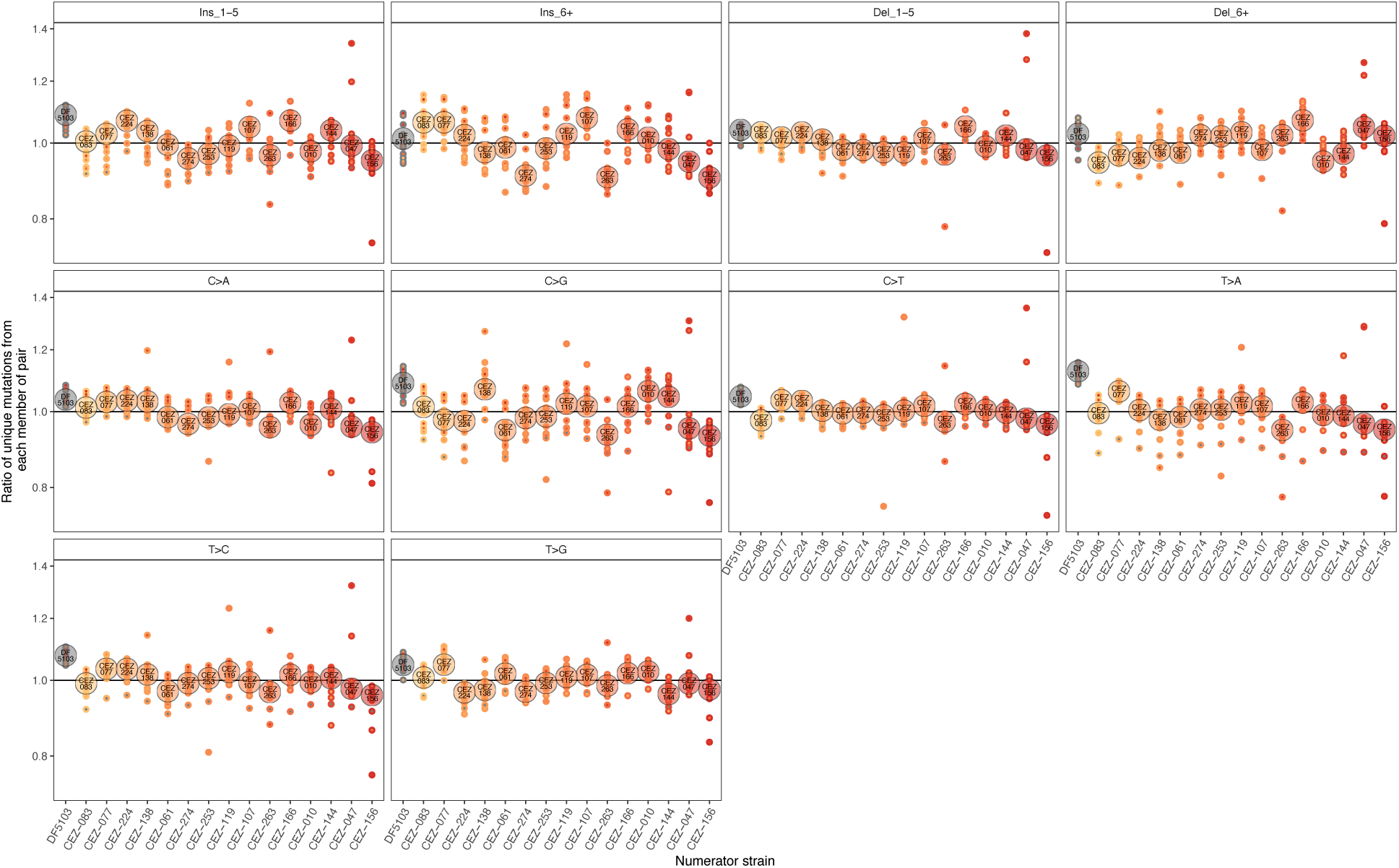
Median (large circles) and pairwise (small circles) relative mutation acquisition between strains, subdivided by mutation type. P-values for glms (Gaussian model) testing association of average relative mutation acquisition to radioactivity are 0.7108 (insertions between 1 and 5 bp), 0.0542 (insertions 6 bp and larger), 0.4862 (deletions between 1 and 5 bp), 0.0906 (deletions 6 bp and larger), 0.0067 (C>A), 0.5610 (C>G), 0.3932 (C>T), 0.1728 (T>A), 0.1039 (T>C), 0.3089 (T>G). Bonferroni corrected significance cutoff = 0.005, which none of these tests surpass. Mantel tests comparing fold change in ambient radiation to relative mutation acquisition between each pair of strains produced a p-value of 1 for all mutation types.

**Supplementary Figure 11.**
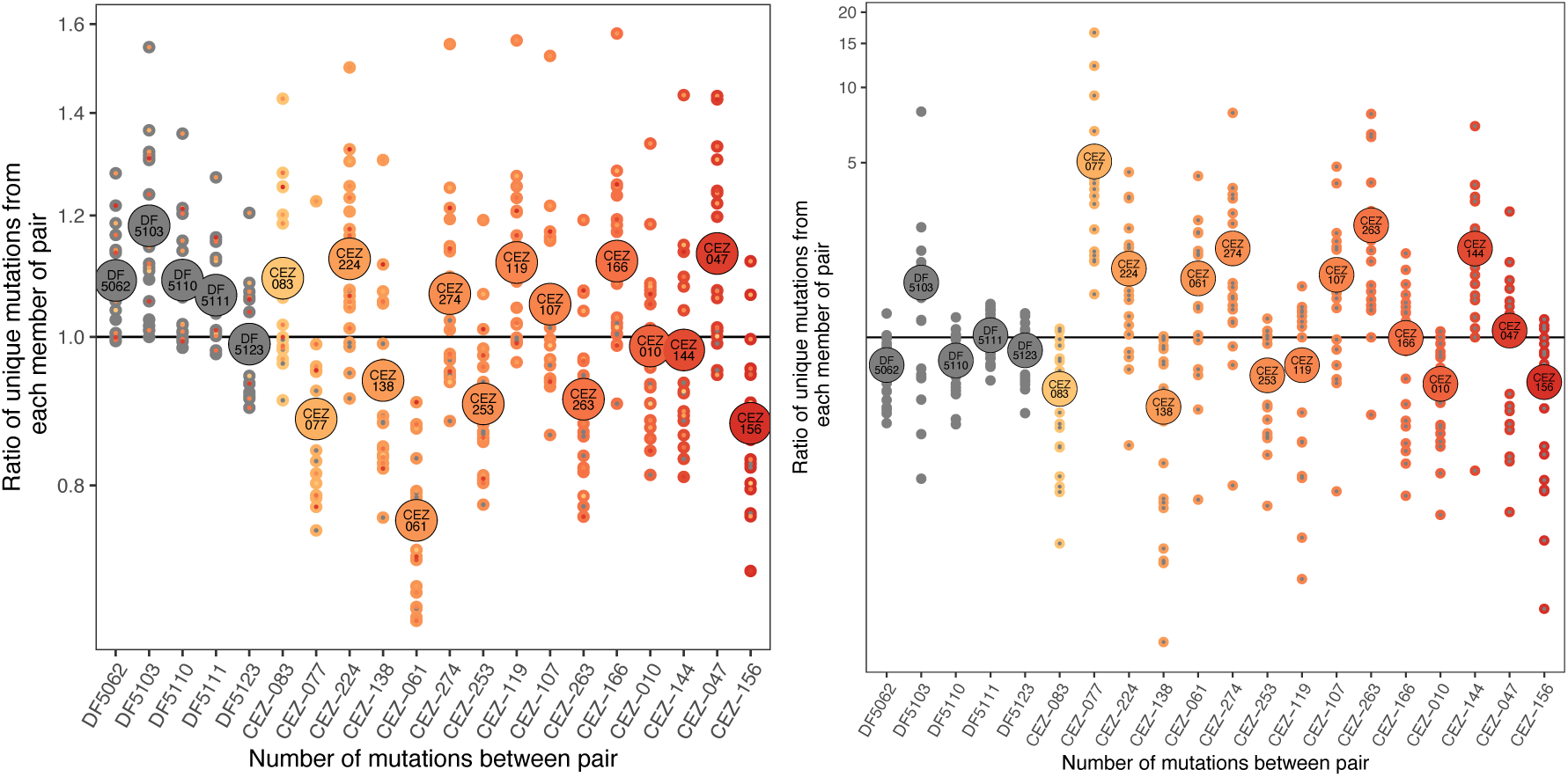
Relative mutation acquisition values for each strain compared to all others, including non-Chornobyl strains*. O. sp. 3 strain* JU75 was used as an outgroup. Small circles represent pairwise comparisons, with the outer color representing the ambient radioactivity of the numerator strain, and the inner color representing the ambient radiation of the denominator strain for each ratio. Large circles represent the average for all comparisons with a common numerator strain. Left: nuclear genome. Right: mitochondrial genome.

**Supplementary Figure 12.**
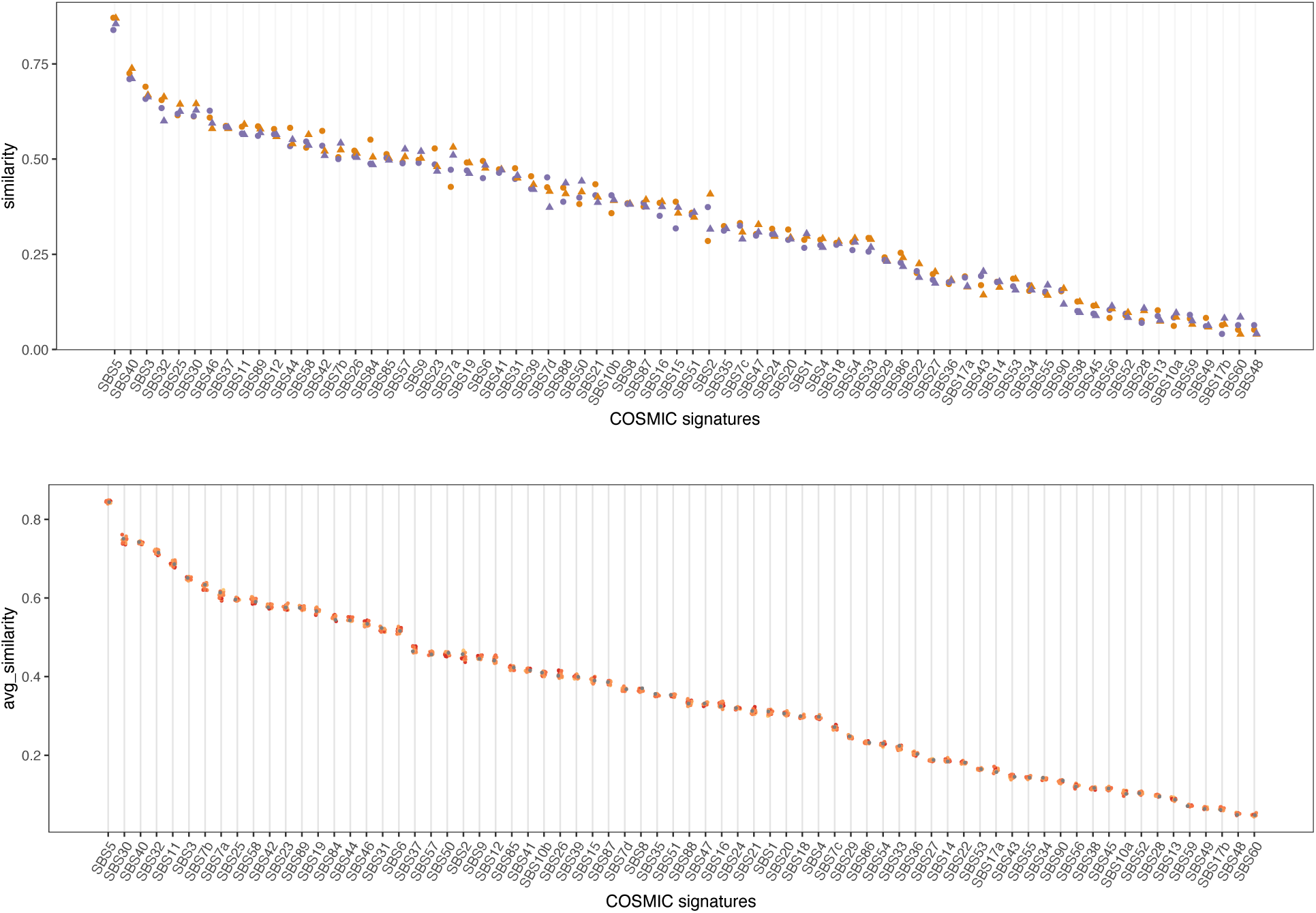
Similarities of *O. tipulae* mutation spectra to COSMIC Single Base Signatures (*28*). Top: Similarities to COSMIC signatures for CEZ-047 compared to CEZ-156 and CEZ-156 compared to CEZ-047 (circles) and CEZ-119 compared to CEZ-253 and CEZ-253 to CEZ-119 (triangles). In both closely related pairs, the one with the higher relative mutation acquisition is in orange (CEZ-047 and CEZ-119) and the lower relative mutation acquisition is in purple (CEZ-156 and CEZ-253). Bottom: Mutation signature were generated from relative mutation acquisition counts for each strain compared to each strain, and then averaged among all comparisons with the same numerator. These averages are plotted, with dots colored according to ambient radioactivity at site where numerator strain was collected. Independent Mantel tests for each COSMIC signature, test whether similarity corresponds with radiation, yielded a p-value of 1 in all cases.

**Supplementary Figure 13:**
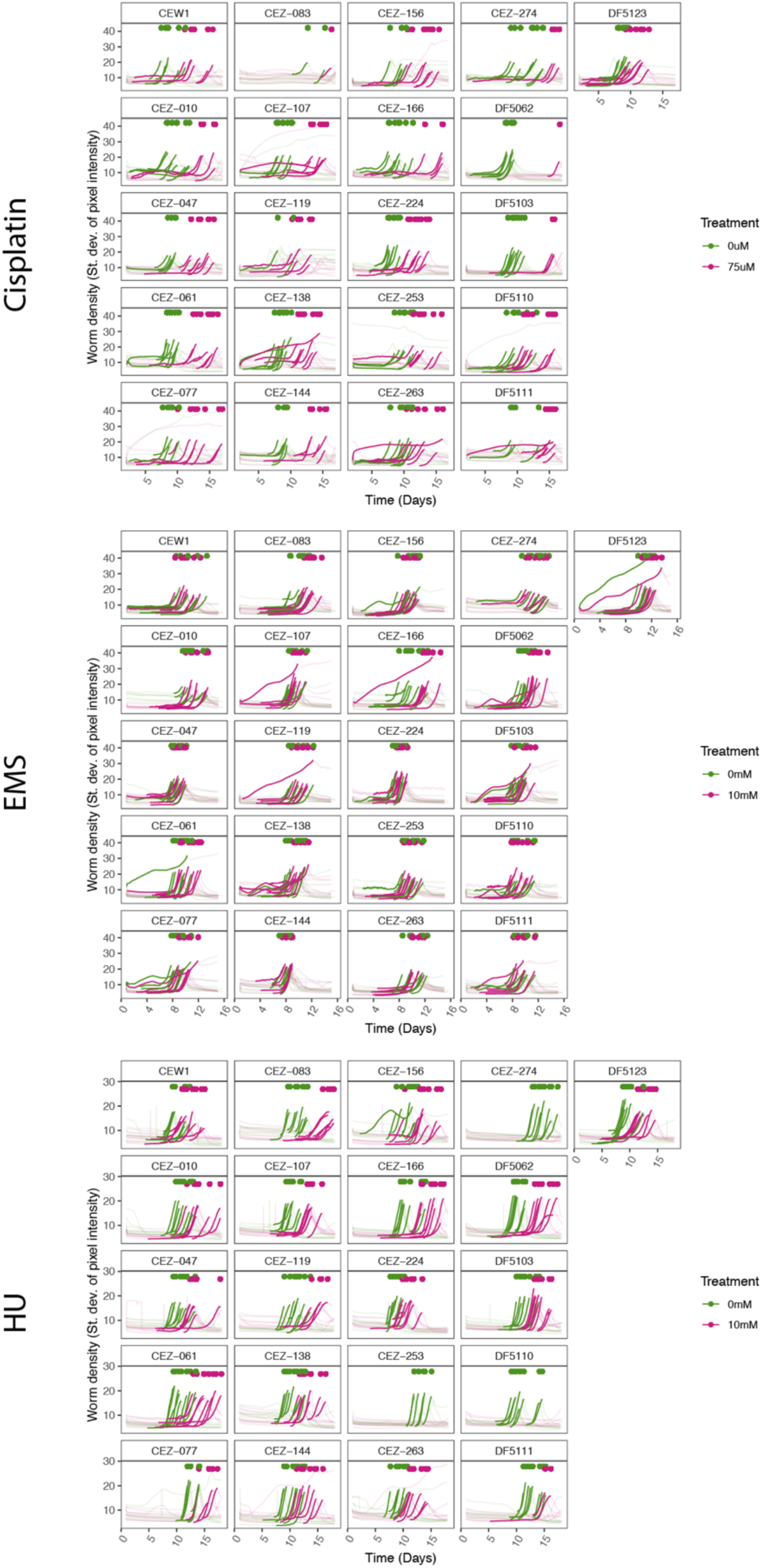
Population growth traces for Figure 5. Images were collected once per hour, and analyzed for the standard deviation of their pixel intensities. Plates with no worms (either before growth or after starvation) have relatively uniform pixel intensity (the solid color of an empty NGMA plate) and so std. dev. of pixel intensity is low. As worm populations grow, pixel intensity becomes bimodel with light pixels from the plate and dark pixels from the worms. Larger the worm populations create higher standard deviations of pixel intensities, as an increasing number of dark worm pixels widens the distribution of the pixel intensities in each image. Because the datapoint of interest is the time at which populations exhaust their food, worm density traces have been grayed out except for the part of the curve leading up to food depletion, to make the endpoint easier to see.

**Supplementary Figure 14:**
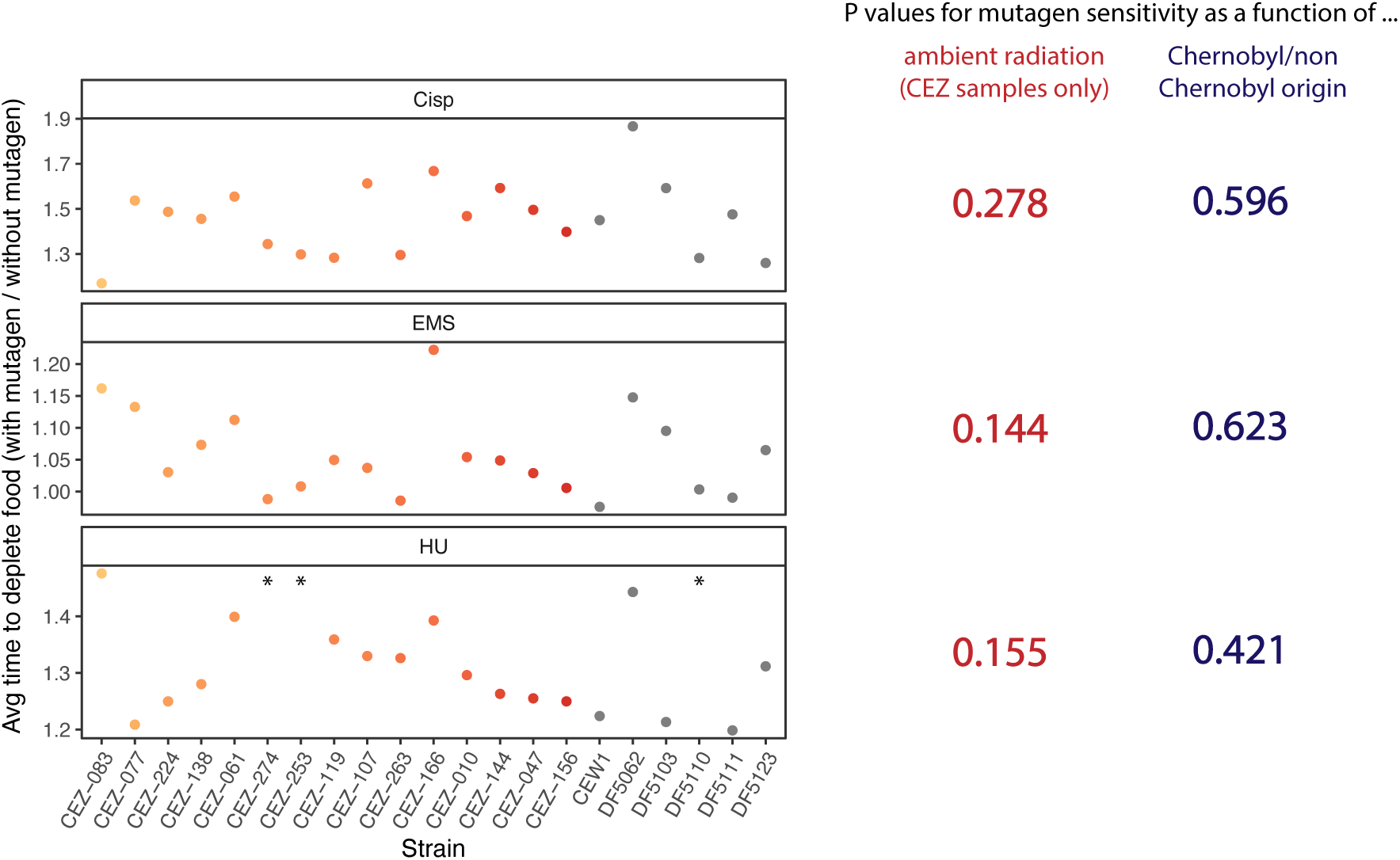
Variation in mutagen sensitivity cannot be explained by ambient radiation at site of collection, or by Chornobyl vs non-Chornobyl sites of collection. P values calculated using glm(family="Gamma") in R (*42*). Asterisks represent samples with no ratio value because no hydroxyurea-treated populations depleted all the available food before the end of the experiment.

**Supplementary Figure 15:**
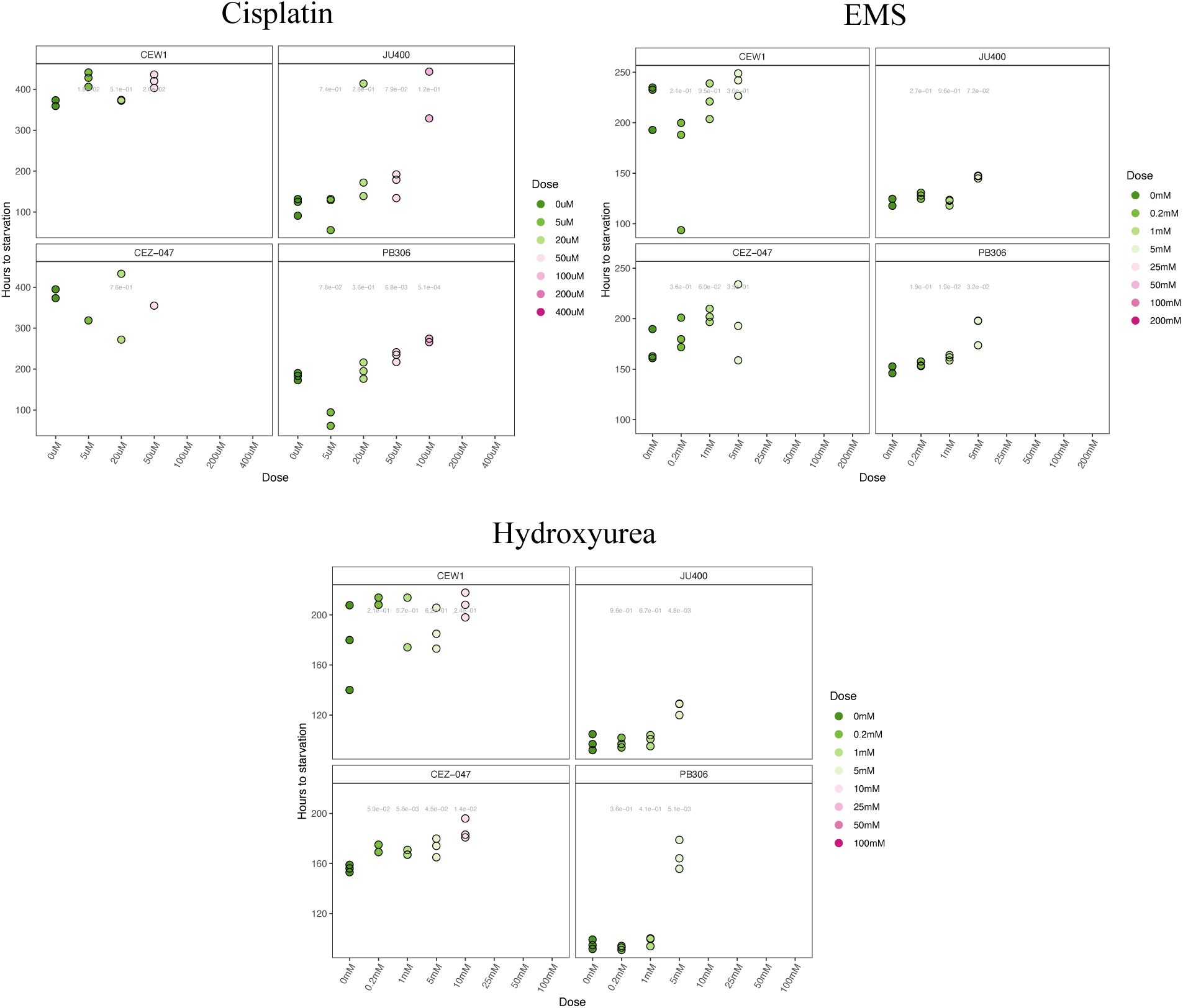
*Oscheius tipulae* (CEW1 and CEZ-047) and *Caenorhabditis elegans* (JU400 and PB306) sensitivities to cisplatin, EMS, and HU. Points missing for replicates that were either contaminated or never grew enough to consume all the food on the plate. *C. elegans* but not *O. tipulae* grew to carrying capacity on 100 uM cisplatin, while *O. tipulae* but not *C. elegans* grew to carrying capacity on 10 mM hydroxyurea.

## Notes

### Competing Interest Statement

The authors have declared no competing interest.

